# BOSE: A Bayesian Order Statistics-Based Estimator for Recovering the Sample Mean and Standard Deviation from the Median, Range, and/or Interquartile Range

**DOI:** 10.64898/2026.06.26.734829

**Authors:** Wenqisi Pan, Zeyu Lu, Wei Jiang, Johan Lim, Lin Xu, Xinlei Wang

**Author notes:** Correspondence to Xinlei Wang. Co-first authors: Wenqisi Pan and Zeyu Lu.

## Abstract

In meta-analyses of continuous outcomes, the sample mean and standard deviation (SD) are essential for synthesizing effect sizes across studies. However, clinical studies frequently report alternative summary statistics, such as the median, quartiles, and range. To enable inclusion of such studies, various methods have been proposed to estimate the sample mean and SD from these reported summaries. We propose the Bayesian Order Statistics-based Estimator (BOSE), which leverages the joint likelihood of observed order statistics together with weakly informative priors to obtain the full posterior distribution for the mean and SD without relying on computationally intensive iterative procedures such as Markov chain Monte Carlo algorithms. Our numerical studies demonstrate that BOSE performs competitively with existing approaches in estimating the mean, while providing comparable or improved SD estimation across the evaluated scenarios, with gains particularly evident in small-sample settings. Under non-normal distributions including skewed, heavy-tailed, and bimodal settings with mild or moderate deviations from normality, BOSE remains robust and stable, whereas methods specifically designed for skewed distributions may become unstable or even inapplicable. Beyond point estimation, BOSE naturally provides empirically validated posterior credible intervals, enabling researchers to formally quantify uncertainty for study-level estimates and make reliable, evidence-based decisions in meta-analytic research synthesis. A publicly accessible web application implementing BOSE and competing methods is also provided to facilitate practical use in meta-analytic research.

## 1 Introduction

Meta-analysis has emerged as a fundamental statistical technique in evidence-based medicine^1^ for synthesizing results across multiple studies to obtain more reliable estimates of treatment effects. For continuous outcomes, standard meta-analytic approaches require each component study to provide study-specific summary statistics, typically the sample mean and standard deviation (SD), to enable proper pooling of data. However, a practical challenge arises when individual studies report heterogeneous summary statistics: while some studies provide the sample mean and SD, others report alternative measures such as the median, quartiles, and range, even for normally distributed data, where such reporting choices may reflect journal preferences, formatting constraints, or analytical traditions within specific medical fields^2^. This inconsistency in reported statistics creates a methodological dilemma for researchers, who must either exclude studies that are missing the required sample mean and SD or develop methods to estimate them from the available summary statistics, thereby maintaining the integrity of the evidence synthesis.

To address this issue, researchers have developed various statistical methods to estimate the sample mean and SD from other commonly reported statistics. Such methods often use the five-number summary (including the sample median *m*, the first and third quartiles *q*_1_ and *q*_3_, and the minimum and maximum values *a* and *b*) or even three-number subsets such as {*a, m, b*} and {*q*_1_, *m, q*_3_}. Typically, this is done along with the sample size *n*, enabling meta-analyses without excluding potentially studies. While these estimation approaches may not perform as well as directly reported sample means and SDs, they provide a practical solution for harmonizing heterogeneous data formats across studies.

Early approaches required no distributional assumptions. Hozo, Djulbegovic, and Hozo^3^ proposed a straightforward method that used the median and extrema {*a, m, b*} along with the sample size *n* to estimate the sample mean and variance. This method was later extended by Bland^4^ to accommodate situations where the first and third quartiles {*q*_1_, *q*_3_} were also available. Subsequent methods introduced more sophisticated formulations under the assumption of normality. Wan et al.^2^ identified a key shortcoming of the approach proposed by Hozo, Djulbegovic, and Hozo^3^: the estimators treated sample size *n* merely as a threshold for selecting fixed formulas, without incorporating its actual value into the estimation, resulting in biased and non-smooth estimates. To address this, Wan et al.^2^ developed improved formulas that fully incorporated the sample size information, significantly enhancing the estimation of the sample SD. They also extended the method to situations where only the median and the first and third quartiles {*q*_1_, *m, q*_3_} are available. With a similar idea, Luo et al.^5^ introduced new estimators for the sample mean that effectively reduce estimation error. Shi et al.^6^ restricted their focus on SD estimation based on the full five-number summary {*a, q*_1_, *m, q*_3_, *b*} and proposed a smoothly weighted estimator to further improve the SD estimator of Wan et al.^2^. More recently, Yang, Hutson, and Wang^7^ introduced the Best Linear Unbiased Estimator (BLUE) for estimating both the sample mean and SD. Among formula-based methods, the approaches developed by Luo et al.^5^ and Wan et al.^2^ are regarded as the most accurate for estimating the sample mean and SD, respectively^8^.

Building on these foundational works, more recent efforts have shifted toward addressing skewed or other non-normally distributed data. McGrath et al.^8^ proposed the quantile estimation (QE) and Box–Cox (BC) methods, and Cai, Zhou, and Pan^9^ further developed the maximum likelihood for non-normal distributions (MLN) method. The QE approach pre-specifies several candidate parametric families, including the normal, log-normal, gamma, beta, and Weibull distributions, and estimates their parameters by minimizing the distance between observed and theoretical quantiles, selecting the best-fitting distribution for downstream estimation. In contrast, the BC and MLN methods apply the Box–Cox transformation to the reported summary statistics, with MLN further employing a likelihood-based framework, constructed from the joint distribution of order statistics, to jointly estimate the transformation and distributional parameters.

Despite these methodological advances, existing approaches retain important limitations. Skewness-based methods such as BC, MLN, and QE may underperform when data do not substantially deviate from normality, and often rely on transformations or candidate distributions requiring strictly positive support, limiting their applicability to data containing zero or negative values. In such settings, these methods may experience numerical instability, convergence failures, or forced fallback to normality-based approximations, thereby losing their ability to effectively capture skewness, heavy tails, or multimodality. Conversely, while normality-based estimators are generally more stable and broadly applicable, they often exhibit reduced accuracy, particularly for SD estimation and in small-sample regimes. Furthermore, the vast majority of existing methods yield only point estimates, with limited consideration given to uncertainty quantification. Although the approach by Yang, Hutson, and Wang^7^ provides variance estimators that enable asymptotic confidence intervals for meta-analytic pooling, the empirical coverage properties of these study-level intervals had not previously been formally evaluated. This highlights a gap in the literature: a need for robust estimators that maintain high efficiency across a range of distributional shapes and sample sizes while concurrently providing well-calibrated uncertainty quantification for individual studies. Specifically, methods should offer interval estimates with empirically validated coverage properties to facilitate more reliable inference, particularly in downstream applications like meta-analysis.

To bridge this gap, we propose a Bayesian Order Statistics-based Estimator (BOSE), which leverages the theoretical properties of order statistics to estimate sample mean and SD from summary statistics reported in primary studies. BOSE approximates the posterior distribution by discretizing the parameter space over a predefined conservative grid, allowing for direct computation of posterior summaries such as point estimates and credible intervals from the discrete empirical CDFs. This approach avoids reliance on asymptotic approximations and does not require computationally intensive, iterative sampling algorithms such as the Gibbs sampler and Metropolis-Hastings algorithm. By construction, BOSE is applicable to any combination of available order statistics, offering a natural extension beyond the scenarios considered in this work. Compared to MLN that also employs a likelihood function of order statistics, a major contribution of BOSE is its capacity for uncertainty quantification, yielding full posterior distributions and empirically calibrated credible intervals, rather than producing singular point estimates. This is useful for meta-analytic applications, where the precision and variability of individual study estimates inform the reliability of downstream synthesis.

Through evaluation, we demonstrate that BOSE achieves competitive or improved performance across a wide range of distributional settings, including normal distributions, mildly to moderately skewed distributions, heavy-tailed distributions, and bimodal mixtures, under varying sample sizes and summary statistics combinations. While methods specifically designed for highly skewed data (e.g., BC, QE, MLN) are, in principle, suited to such settings, our simulation results indicate that their performance can deteriorate under more subtle departures from normality, and in small samples, this instability can persist under more pronounced skewness. In many practical applications, the extent of non-normality is difficult to infer from reported summary statistics alone: pronounced skewness may be reflected by substantial asymmetry between the distances of the maximum and minimum from the median, but more commonly, only mild or moderate asymmetry is observed. Such deviations may arise either from genuinely skewed underlying distributions or from sampling variability, rendering reliable discrimination challenging. In such non-extreme regimes, our proposed method exhibits stable and robust performance, making it a practical default choice.

The remainder of this paper is organized as follows. Section 2 presents the methodological framework, introducing the proposed Bayesian estimation method. Section 3 describes simulation studies designed to evaluate and compare the performance of all methods across various scenarios and sample sizes. Section 4 demonstrates the practical applications of these methods through real-world data analysis, illustrating their implementation in actual meta-analytic contexts. Finally, Section 5 concludes with a discussion of key findings and methodological implications.

## 2 Method

### 2.1 The Data Model

Consider an individual study included in a meta-analysis, based on a sample of size *n*, {*X*_1_, *X*_2_, …, *X*_*n*_}, assumed to be i.i.d. random variables from a continuous parametric distribution with probability density function (PDF) *f*_*θ*_ (*x*) and cumulative distribution function (CDF) *F*_*θ*_ (*x*), where *θ* denotes the parameter vector. Let *X*_(1)_ ≤ *X*_(2)_ ≤ … ≤ *X*_(*n*)_ denote the corresponding order statistics. A meta-analysis typically requires the sample mean and the sample SD of each component study to estimate the overall (standardized) effect size. However, as mentioned in the introduction, a significant challenge in meta-analysis is that some component studies report alternative summary statistics rather than sample mean 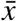 and SD *s* of {*X*_1_, *X*_2_, …, *X*_*n*_}. These alternative summary statistics (such as median, quartiles, range) are derived from a subset of the order statistics from the original sample. We therefore assume that, for a given study, a subset of *k* order statistics 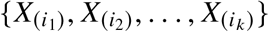with 1 ≤ *i*_1_ < *i*_2_ < … < *i*_*k*_ ≤ *n* is observed instead of the sample mean and SD. Based on the theory of order statistics^10^, the joint density function of this subset is given by:

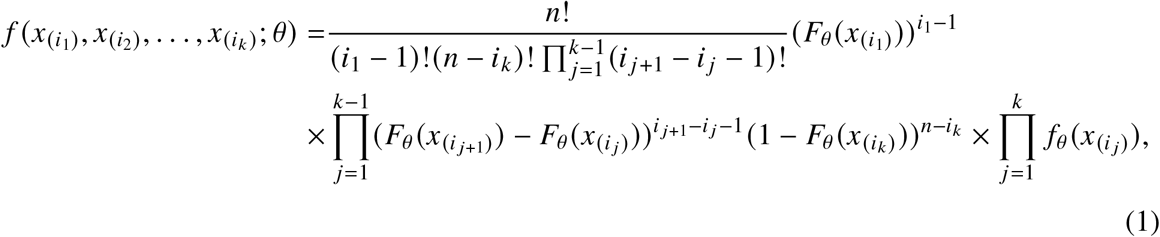

where 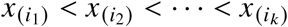.

In practice, the specific order statistics entering the likelihood depend on the definition of the sample quantile employed. Hyndman and Fan^11^ summarized nine existing definitions, commonly referred to as Types 1-9, which can be broadly classified into two groups: discontinuous definitions (Types 1–3), which select an observed order statistic according to a predetermined index rule (e.g., ceiling), and continuous definitions (Types 4–9), which are based on linear interpolation between order statistics. Let ⌈·⌉ denote the ceiling function. In our method, we adopt the Type 1 quantile definition throughout, under which *i* = ⌈*np*⌉, where *n* is the sample size, *p* ∈ (0, 1) is the probability level of a quantile, and *X*_(*i*)_ denotes the *i*th order statistic, which gives a direct correspondence between quantiles and order statistics.

In meta-analysis, for studies that report alternative summary statistics rather than 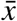 and *s*, three scenarios commonly arise, namely *S*_1_ = {*a, m, b*; *n*}, *S*_2_ = {*q*_1_, *m, q*_3_; *n*}, and the complete five-number summary *S*_3_ = {*a, q*_1_, *m, q*_3_, *b*; *n*}. Under the Type 1 quantile definition, the corresponding order statistic indices for a sample with size *n* are:

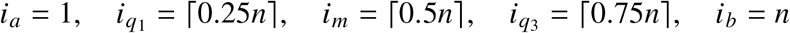

Ignoring the normalizing constant in Equation (1), the likelihood functions in the three scenarios can be expressed as:

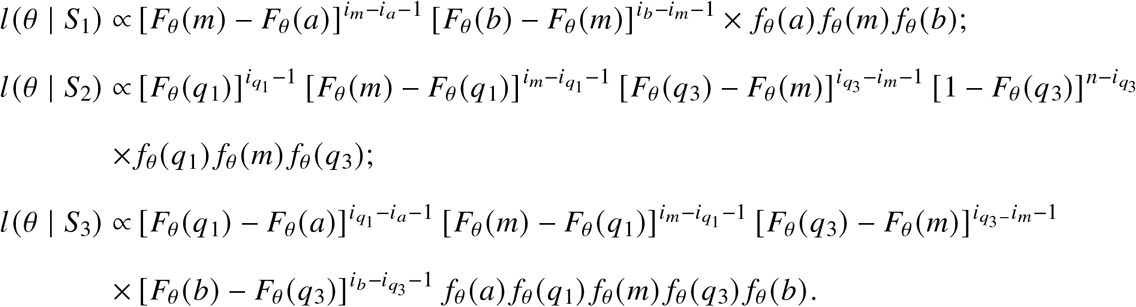

Although these likelihood functions are applicable to any continuous distribution, we proceed with normal distributions, as is typical in meta-analysis for continuous outcomes.

### 2.2 Prior Specification

Let *ϕ*(·) and Φ(·) denote the probability density function (pdf) and the cumulative distribution function (cdf) of *N* (0, 1), respectively. Then, for the normal model *X*_*i*_ ~ *N* (*µ, σ*^2^), where 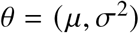, we have 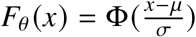 and 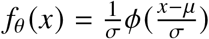. We adopt weakly informative priors for *θ* that balance objectivity with computational stability. The use of weakly informative priors, rather than improper reference priors, was recommended by Gelman^12^ and has been successfully adopted in many Bayesian applications such as Jia et al.^13^ and Lu, Xu, and Wang^14^. These priors ensure proper posterior distributions while still allowing the data to primarily drive the inference. Specifically, we consider *a priori* independent priors on *µ* and *σ*^2^:

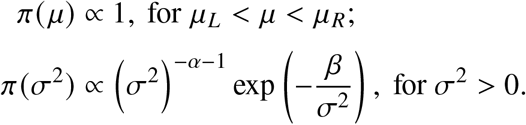

That is, for the location parameter *µ*, we assign a uniform prior over the interval (*µ*_*L*_, *µ*_*R*_), where *µ*_*L*_ and *µ*_*R*_ are the lower and upper bounds delineating a plausible range. We further set 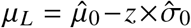 and 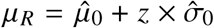, with 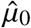 derived from the method proposed by Luo et al.^5^ and 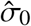 obtained following the approach of Wan et al.^2^; the scaling factor *z* controls the width of the interval and can be chosen to define a broad range for *µ* (e.g., *z* = 3, 5, 10). Because the uniform prior bounds are calculated from the same reported summary statistics used in the likelihood, BOSE adopts an empirical-Bayes formulation. The interval is deliberately broad and is motivated primarily by computational efficiency. Within this interval, the prior density is constant; its influence on the posterior arises through truncation at the boundaries. This influence is expected to be small when the posterior is concentrated well inside the interval, consistent with the minimal variation observed in our sensitivity analysis across *z* = 3, 5, and 10 (Section 3.1; Supplementary Figure S1). Posterior distributions and credible intervals are computed conditional on these selected bounds. For the variance parameter *σ*^2^, we specify an inverse-gamma prior IG(*α, β*), where *α* and *β* are selected to be small positive numbers (e.g., 0.01), approximating a diffuse prior with minimal information, as discussed by Spiegelhalter, Abrams, and Myles^15^.

### 2.3 Posterior Computation and Inference

We derive the (unnormalized) joint posterior distribution of (*µ, σ*^2^) for the three scenarios *S*_1_–*S*_3_:

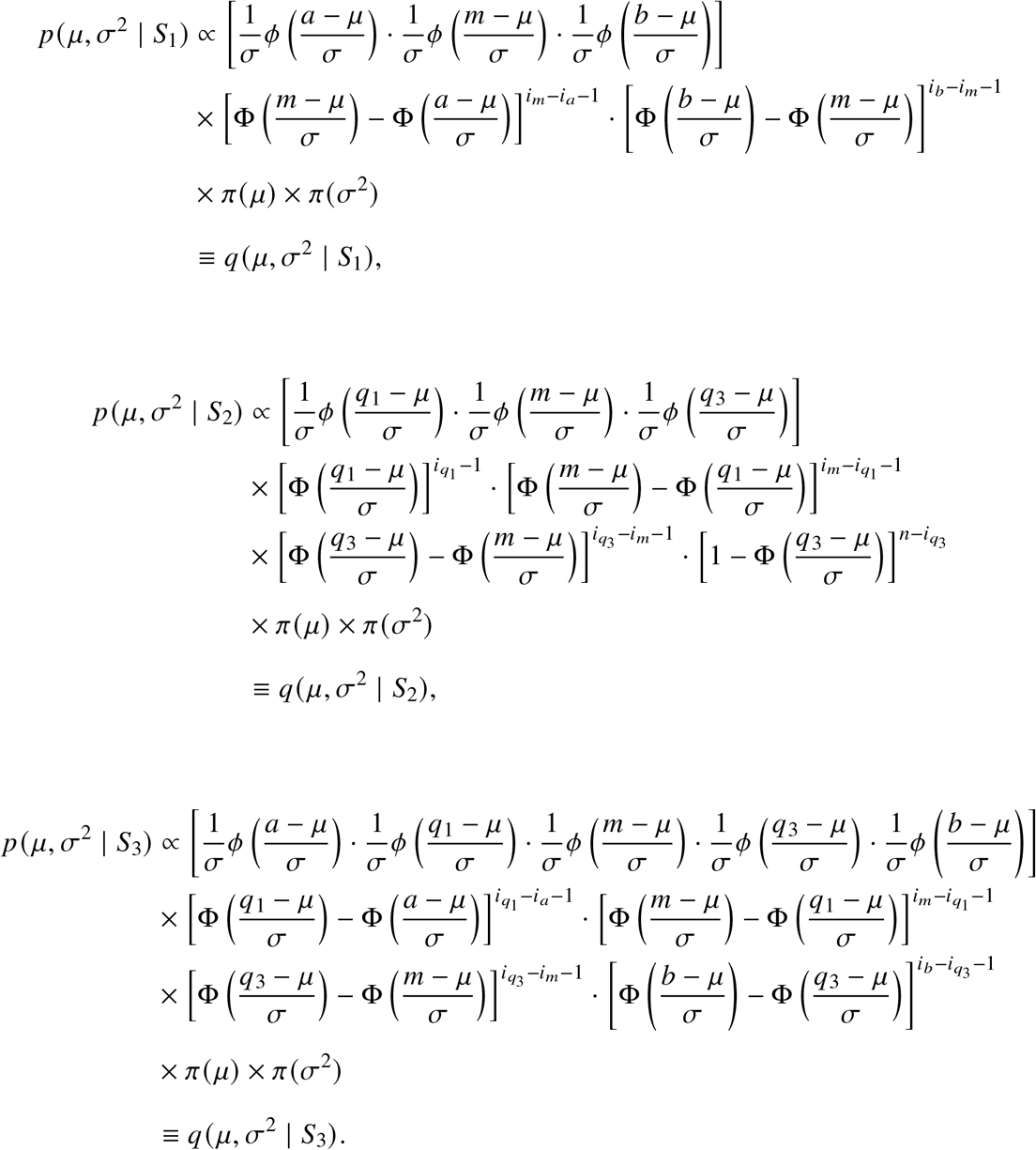

However, neither the joint posterior distributions nor their full conditionals belong to any known parametric family, rendering direct sampling infeasible and making Gibbs sampling unsuitable. We instead approximate the joint posterior by evaluating the (unnormalized) log-posterior over a two-dimensional grid spanning a plausible range of values for *µ* and *σ*. This is a technique that has long been used for low-dimensional Bayesian computation^16^. Given that our setting involves only two parameters, this grid-based approximation is computationally efficient while providing sufficient accuracy. Specifically, we employ a two-stage adaptive strategy: (i) A coarse grid of size *G*_*c*_ × *G*_*c*_, evenly spaced, is used to scan an initial region *R*_0_, defined as 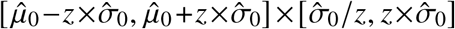, to identify a region of interest (ROI) containing at least 99% of the posterior mass. At each grid point (*µ*_*i*_, *σ*_*j*_), the log (unnormalized) posterior for *S*_*l*_ (i.e., log *q* (*µ, σ*^2^ | *S*_*l*_)) is computed. For numerical stability, these log-*q*values are shifted by subtracting their maximum value and then exponentiated to obtain the weights. Grid points are then ranked by these posterior weights, and weights are accumulated until reaching 99% of the total mass; the ROI is then defined as the bounding rectangle of these high-density points, denoted by [*µ*_*min*_, *µ*_*max*_] × [*σ*_*min*_, *σ*_*max*_]. (ii) A finer grid of size *G* _*f*_ × *G* _*f*_ (typically *G* _*f*_ = 2*G*_*c*_), evenly spaced, is constructed over the ROI expanded by a 10% padding in each dimension. That is, for each parameter, 10% of the ROI width is added to both the lower and upper bounds. This refinement concentrates computational effort in regions of high posterior density while maintaining sufficient coverage.

We derive point estimates and quantify uncertainty using the marginal posterior distributions. Our fine-grid approximation allows us to seamlessly compute the normalized marginal probabilities. By marginalizing over *σ*^2^, the marginal probability for each *µ*_*i*_ can be approximated by:

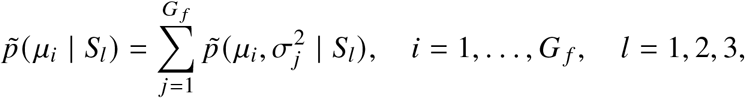

where 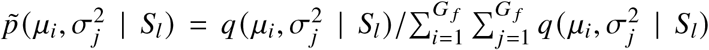 is the approximate posterior at the (*i, j*) grid point. The marginal probabilities for 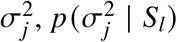, can be similarly approximated by 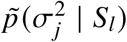 by summing across the rows.

For the point estimate of the location parameter, we utilize the posterior mean: 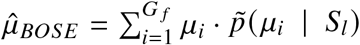. Due to the potential skewness of the marginal posterior distribution for the scale parameter, we use the posterior median as a robust point estimate for *σ*, denoted as 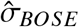. Both the posterior median and the 100(1 − *α*)% central credible intervals (CI) are computed efficiently via cumulative distribution function (CDF) inversion. Let 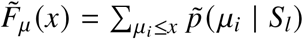 and 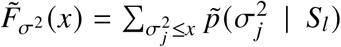 denote the discrete empirical CDFs. The estimates are obtained by finding the corresponding quantiles (e.g., *F*^−1^(0.5) for the median, and *F*^−1^(0.025) and *F*^−1^(0.975) for a 95% CI) using linear interpolation on the grid values. The detailed procedure for Scenario *S*_3_ is presented as an example in Algorithm 1 of the Supplementary Material.

## 3 Simulation

### 3.1 Study Settings

A simulation study is conducted to evaluate the performance of the proposed method BOSE in parameter estimation and uncertainty quantification. We first specify the hyperparameter settings of BOSE used in our simulation. For the inverse-gamma prior IG(*α,β*) on *σ*^2^, we set *α* = *β* = 0.01. For the uniform prior on *µ*, the scaling factor *z* is set to 5. Results from a sensitivity analysis (Supplementary Figure S1), examining all combinations of *z* ∈ {3, 5, 10} and *α* = *β* ∈ {0.1, 0.01, 0.001}, show minimal variation across settings, indicating that BOSE is robust to prior specification. Therefore, we fix *z* = 5 and *α* = *β* = 0.01 in all subsequent analyses. Similarly, for the coarse and fine grid sizes used in the two-stage grid approximation, we fix *G*_*c*_ = 125 and *G* _*f*_ = 250, with a 10% padding applied when constructing the fine grid to avoid truncating posterior mass near the boundary of the coarse-grid region of interest. Results from a sensitivity analysis varying the grid size (four configurations: (*G*_*c*_, *G* _*f*_) = (64, 128), (125, 250), (200, 400), and (256, 512)) and the padding percentage (5%, 10%, 20%), shown in Supplementary Figure S2, confirm that our results are robust to these choices.

We benchmark BOSE against other normality-based methods and skewness-based methods across all simulated scenarios, encompassing simulations under the normality assumption as well as robustness assessments under non-normal distributions. The normality-based competitors include LW, which combines Luo et al.^5^ and Wan et al.^2^ for estimating the sample mean and SD, respectively; BLUE; and the Shi method, which is specifically designed for SD estimation in Scenario *S*_3_. The skewness-based benchmark methods include BC, QE, and MLN.

To assess estimation performance under the normality assumption, we consider two distributions, *N* (50, 17^2^) and *N* (5, 1^2^), first used by Hozo, Djulbegovic, and Hozo^3^ and Bland^4^, respectively, and subsequently adopted in many related studies. Data are generated using Type 1 sample quantiles. We additionally conduct simulations across all nine sample quantile definitions (Types 1–9) under these two normal settings to evaluate the sensitivity of the estimators to the specific quantile definition employed.

To evaluate robustness to departures from normality, we consider non-normal distributions representing three key characteristics: skewness, heavy tails and bimodality. Because BOSE is developed under the normality assumption, our primary assessment focuses on mild to moderate departures, with additional settings used to examine its limits under more pronounced skewness and heavier tails. For skewness, we employ log-normal and gamma distributions with target skewness values *κ* ∈ {0.1, 0.3, 0.5, 0.7, 0.9}, representing the small to moderate skewness of primary interest. The distributional parameters are determined analytically for each target skewness level. Specifically, we simulate data from Log-Normal(5, *σ*^2^) distributions, where *σ*^2^ is obtained by solving the skewness equation 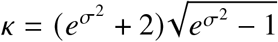, and from Gamma(*α*, 1) distributions, where the shape parameter *α* satisfies 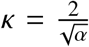. We also consider *κ* ∈ {1, 2, 3} for both families to assess performance under stronger skewness. To assess heavy-tail effects, we employ Student’s *t* distributions with degrees of freedom *df* ∈ {5, 10, 15, 20, 25}. These settings range from the heavier-tailed *t*_5_ distribution to the approximately normal *t*_25_ distribution. We further examine *df* ∈ {3, 4} to assess robustness under more extreme heavy tails. For bimodality assessment, we simulate from the mixture normal distribution *pN* (−1, 1) + (1 − *p*) *N* (1, 1), where the mixing proportion *p* ∈ {0.1, 0.2, 0.3, 0.4, 0.5} controls the degree of component imbalance.

For both normal and non-normal settings, we perform *T* = 1, 000 replicates. For normal settings, sample sizes range from *n* = 5 to 200, similar to those used by Wan et al.^2^ and Yang, Hutson, and Wang^7^. For non-normal settings, we use *n* = 5 to 50 to focus on small-sample performance. In these settings, unstable empirical quantiles and limited information about tail behavior can prevent three- and five-number summaries from adequately capturing distributional asymmetry. These limitations make estimation from order statistics particularly challenging and may accentuate differences between methods. For larger sample sizes under non-normality, one can apply a skewness test to decide whether the study should be excluded^17^ or use skewness-based methods, both of which tend to perform well.

Throughout our simulation, we consider the three data reporting scenarios: *S*_1_ = {*a, m, b*} (minimum, median, maximum), *S*_2_ = {*q*_1_, *m, q*_3_} (1st and 3rd quartiles and median), and *S*_3_ = {*a, q*_1_, *m, q*_3_, *b*} (full five-number summary). Point estimation performance is evaluated using the relative mean squared error (RMSE), following previous work such as Luo et al.^5^ and McGrath et al.^8^, which measures estimation efficiency relative to the corresponding sample statistics. Let 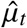 and 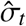 denote the estimated population mean and SD in the *t*-th replicate (*t* = 1, 2, …, *T*), and let 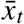 and *s*_*t*_ represent the corresponding sample mean and sample SD. Further, let *µ* and *σ* denote the true population mean and SD, respectively. The RMSEs for mean and SD estimation are defined as:

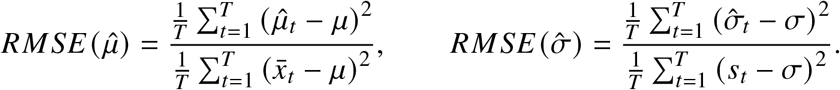

An RMSE value below 1 indicates that the estimator achieves lower mean squared error than the corresponding sample statistic. In practice, for methods estimating the mean and SD from partial quantile information, RMSE values are typically greater than 1, with smaller values indicating better performance. Interval estimation performance is evaluated via comparing the empirical coverage rate (i.e., proportion of the CIs covering the true parameter value) against the nominal level. Throughout this work, calibration refers to this empirical assessment in the simulation settings examined, rather than a theoretical guarantee of nominal coverage across all parameter values or distributions.

Finally, we mention that under normality, all methods perform without computational failures, as the simulated data are strictly positive. However, skewness-based methods (BC and MLN) encounter computational failures when applied to negative values, which arise naturally under heavy-tailed and bimodal distributions. BC and MLN produce errors that are caught and recorded as missing values (“NA” in the method output) in such cases, while QE defaults to fitting a normal distribution. Beyond these explicit failures, all these three methods occasionally produce RMSE values several orders of magnitude beyond the typical range (up to 10^19^ for QE), indicating silent numerical breakdowns. We therefore apply a uniform threshold of 50 to all methods: any replication with RMSE exceeding this value is also set to NA. This cutoff lies well above the maximum RMSE observed for BOSE, LW, BLUE, and Shi (all below 16), and the total exclusion rate is insensitive to the exact choice (varying by fewer than 3 percentage points for thresholds between 20 and 500).

### 3.2 Performance Comparison under Normality

#### 3.2.1 Results of Point Estimation

Figure 1 reports the performance of BOSE and competing methods in estimating the mean (top two rows) and SD (bottom two rows) under data generated from normal distributions. For each method, RMSE was computed across simulation replicates for each individual sample size and then averaged over sample sizes within predefined groups (5–10, 10–30, 30–50, 50–100, and 100–200); the averaged RMSE values are reported in Supplementary Tables S1 (mean) and S2 (SD). The five groups shown here are used only to aid visual presentation. Supplementary Figure S3 reports RMSE for individual sample sizes (displayed at every 10*th* value of n for readability) and shows trends consistent with the grouped averages. Overall, BOSE achieves the lowest or near-lowest RMSE across the settings considered.

**Figure 1:**
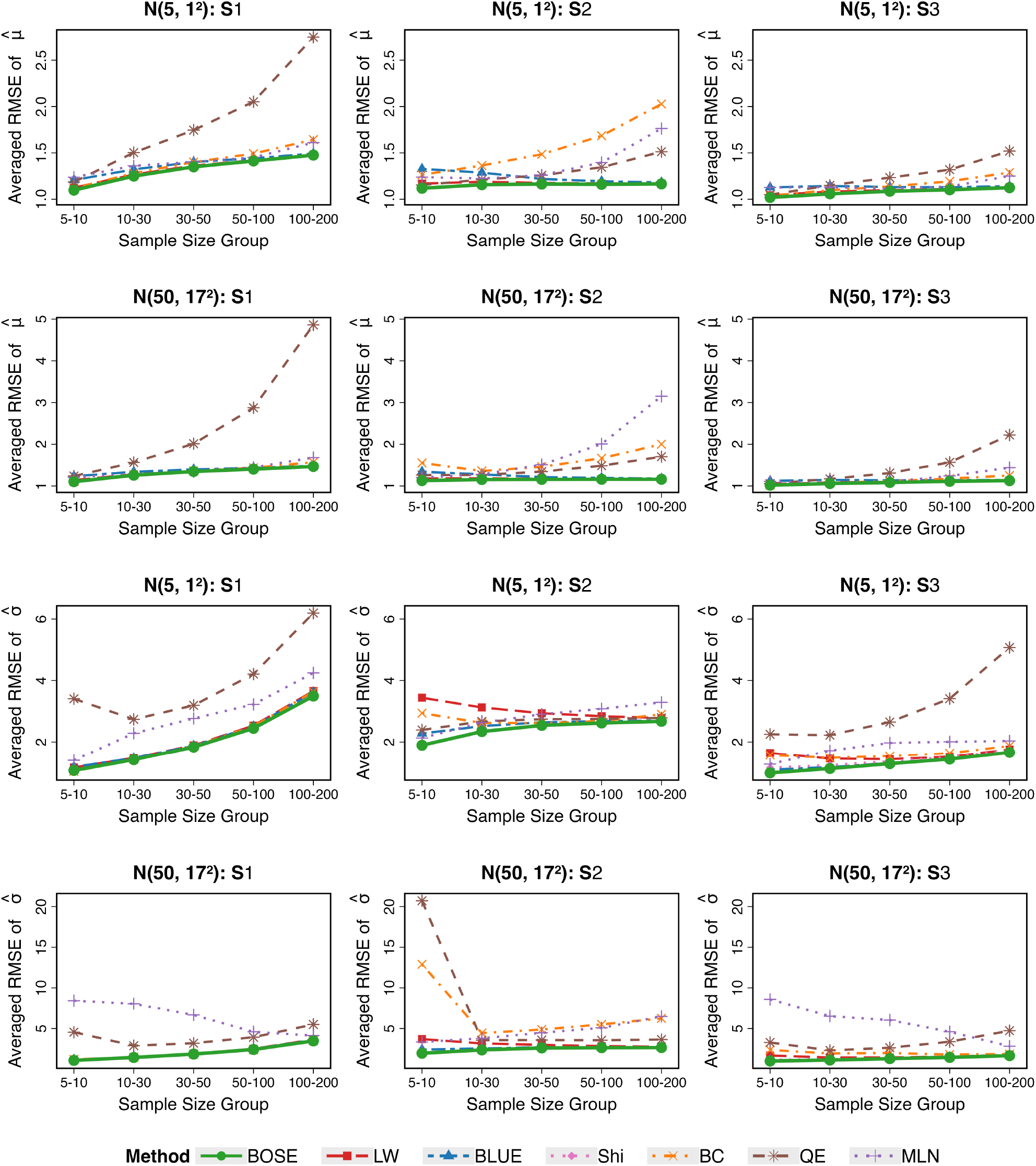
Performance comparison based on averaged RMSE for mean estimation (top two rows) and SD estimation (bottom two rows) using data generated from normal distributions under three common data reporting scenarios *S*_1_ − *S*_3_. For each method, RMSE values are averaged across all sample sizes within each of the five sample-size groups shown (5-10, 10-30, 30-50, 50-100, 100-200). The performance of the proposed BOSE method (solid green line) is compared with the existing methods when applicable across five sample size groups.

As the sample size *n* increases, RMSE tends to increase for most methods, especially for mean estimation. This behavior is expected because the baselines used in the RMSE calculation (i.e., the sample mean and sample SD) become increasingly accurate with larger sample sizes, whereas all considered methods continue to rely only on either five-number or three-number summary statistics, yielding relatively poorer performance.

In general, normality-based methods show better stability across different sample size groups than skewness-based methods, with BOSE appearing to be the least sensitive to changes in sample size. For large *n*, the performance of the two groups can differ substantially. Within the normality-based methods, performance differences become relatively small when *n* is large, with BOSE maintaining a slight advantage. In contrast, BOSE demonstrates a clearer advantage in small-sample settings (*n* ≤ 50), as shown in Figure 2, particularly for SD estimation under Scenarios *S*_2_ and *S*_3_, suggesting that incorporating interquartile information benefits BOSE more substantially than other normality-based methods. This finding is especially relevant in practical meta-analysis applications, where component studies are often constrained by limited sample sizes.

**Figure 2:**
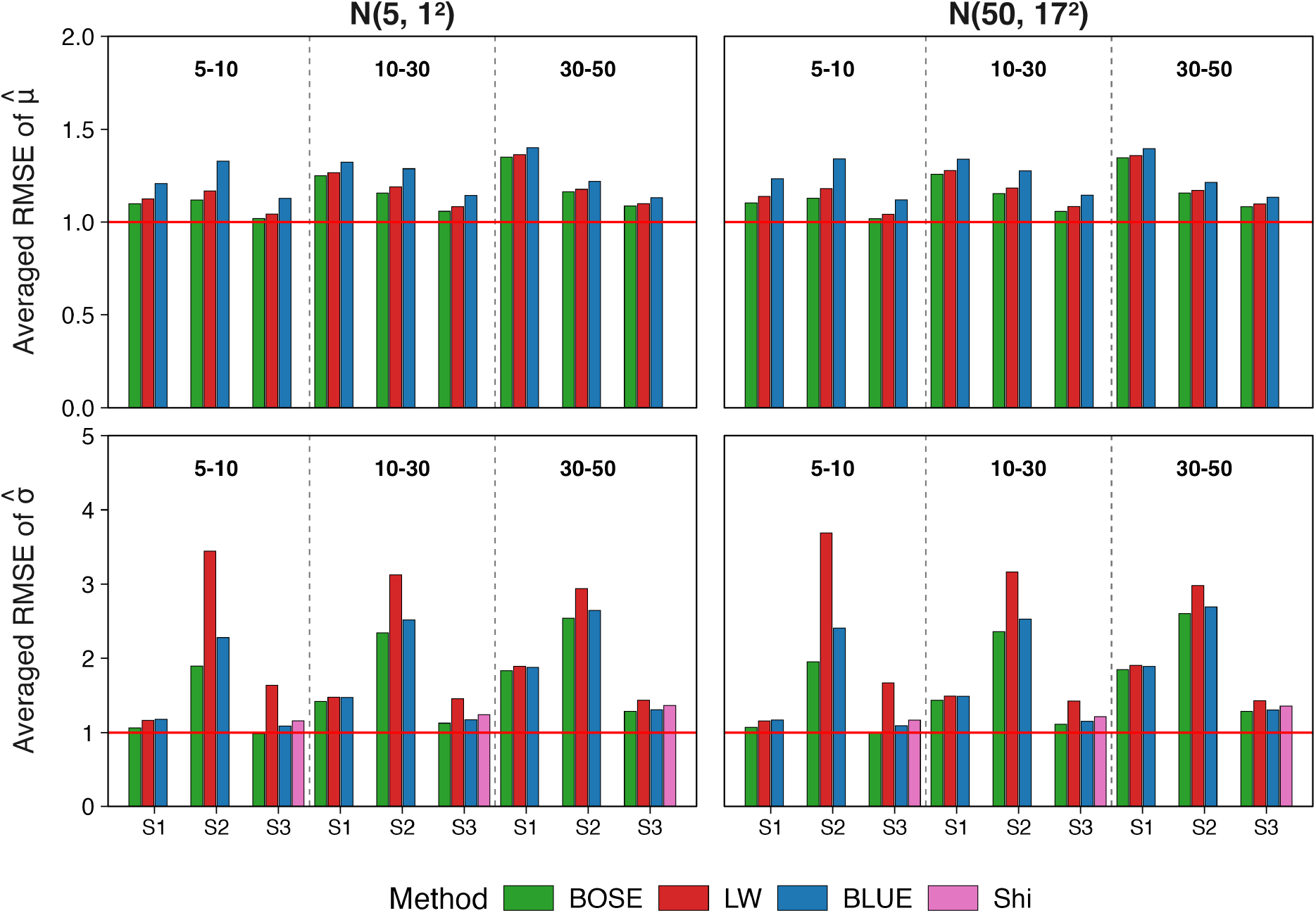
Performance comparison of normality-based methods for small sample size groups where *n* ≤ 50 based on averaged RMSE for mean estimation (top row) and SD estimation (bottom row) using data generated from normal distributions under three common data reporting scenarios *S*_1_ − *S*_3_. For each method, RMSE values are averaged across all sample sizes within each of the three sample-size groups shown (5-10, 10-30, 30-50). The performance of the proposed BOSE method (solid green bar) is compared against other normality-based methods when applicable across three sample size groups.

Across the three reporting scenarios, Scenario *S*_3_ consistently yields lower average RMSE values for most methods, suggesting that incorporating both interquartile and range information improves estimation accuracy. In addition, RMSE values under *S*_2_ are generally lower than those under S1, indicating that the first and third quartiles often provide more informative summaries than the minimum and maximum values.

To further examine whether the observed performance advantage of BOSE is driven by the specific quantile definition used in data generation, we repeat all analyses across sample quantile Types 1 through 9 and report the RMSE values averaged over all sample sizes in Figure 3. BOSE again consistently achieves the lowest averaged RMSE level across all quantile types, scenarios, and distributional settings. Overall, most methods are relatively insensitive to the choice of quantile type, with BOSE among the least sensitive methods, whereas the skewness-based methods BC and MLN tend to be more sensitive. The relative advantage of BOSE becomes particularly evident in settings where all methods exhibit high sensitivity to the choice of quantile type (e.g., Scenario *S*_2_ under *N* (5, 1^2^)). These results demonstrate that the performance of BOSE is insensitive to the quantile types used in data generation, which is desirable in practice because statistical software packages often adopt different default quantile calculation methods (e.g., R uses type 7, SAS uses type 2, and SPSS uses type 6).

**Figure 3:**
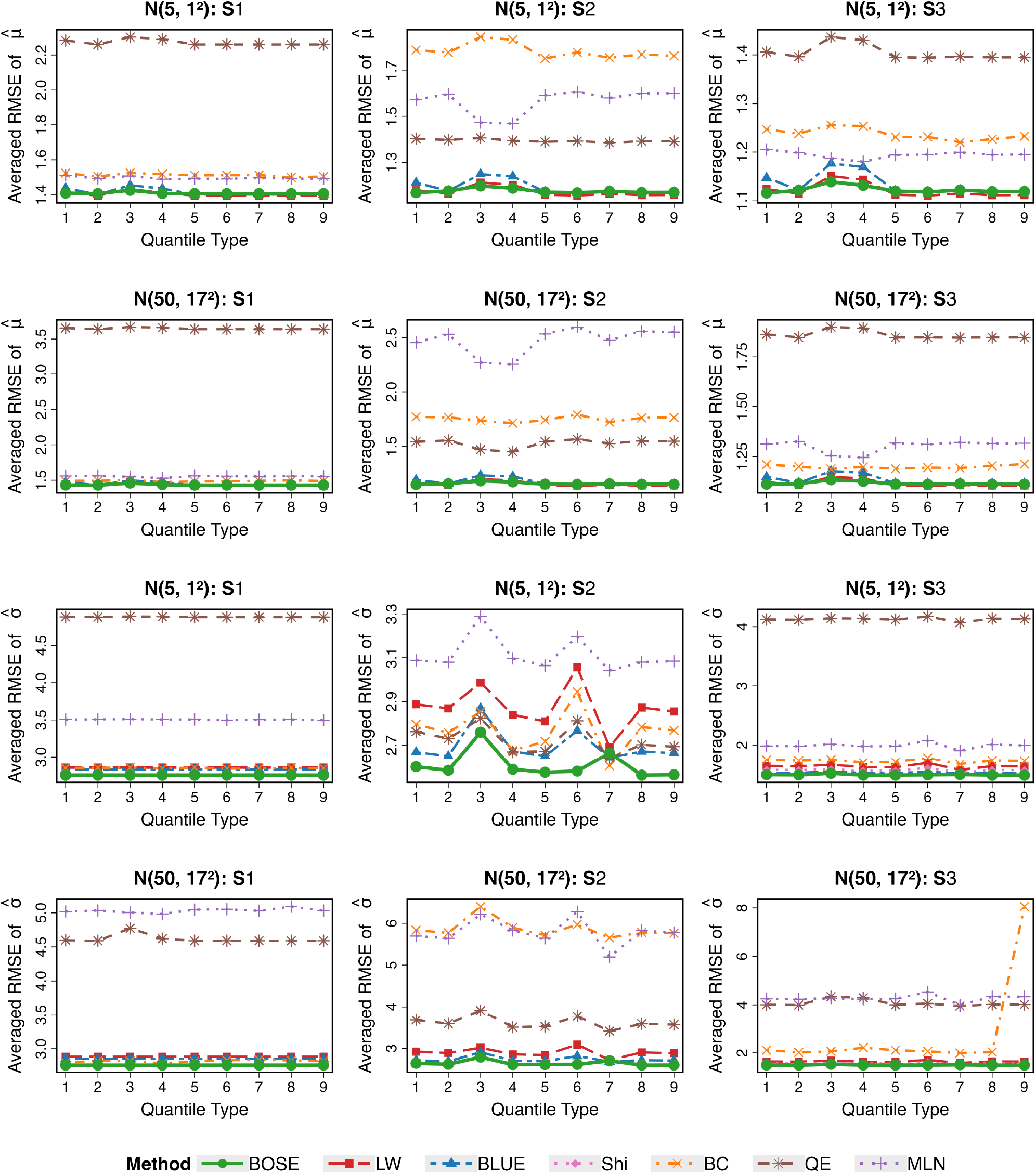
Sensitivity analysis of sample quantile type choices based on averaged RMSE for mean estimation (top two rows) and SD estimation (bottom two rows) using data generated from normal distributions under three common data reporting scenarios *S*_1_ − *S*_3_. The performance of the proposed BOSE method (solid green line) is compared with the existing methods when applicable across nine sample quantile types (Types 1–9).

#### 3.2.2 Results of Interval Estimation

We next evaluate the interval estimation performance of BOSE by examining the empirical coverage probabilities of its credible intervals for both 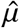 and 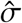 at nominal levels of 95%, 90%, and 80% (Supplementary Figure S4). Across all scenarios and distributional settings, the coverage rates align well with their respective nominal levels, and the narrow interquartile ranges indicate high consistency across simulation replicates.

Figure 4 compares the 95% coverage rates of BOSE and BLUE, the only competing method that produces confidence intervals, stratified by the same five sample-size groups used in Figure 1 (5– 10, 10–30, 30–50, 50–100, and 100–200). This stratification reveals that the coverage advantage of BOSE over BLUE is concentrated in small samples: in the 5–10 and 10–30 groups, BLUE exhibits systematically lower median coverage rates, wider interquartile ranges, and a longer tail of outliers that frequently fall below 0.85, particularly under Scenario *S*_2_, while BOSE remains closely centered near the nominal 0.95 level. Supplementary Figure S5 shows this pattern at the level of individual sample sizes, confirming a smooth, continuous improvement in both methods’ coverage as *n* increases, with BOSE consistently closer to the nominal level than BLUE at small *n*. For sample sizes of 30 or more, coverage for both methods converges closely to the nominal level, and the difference between BOSE and BLUE becomes negligible. This pattern is consistent across both distributional settings and all three data reporting scenarios, and is consistent with the limitations of asymptotic approximations, which rely on large-sample assumptions that may not hold when the sample size is small or only limited summary statistics are available.

**Figure 4:**
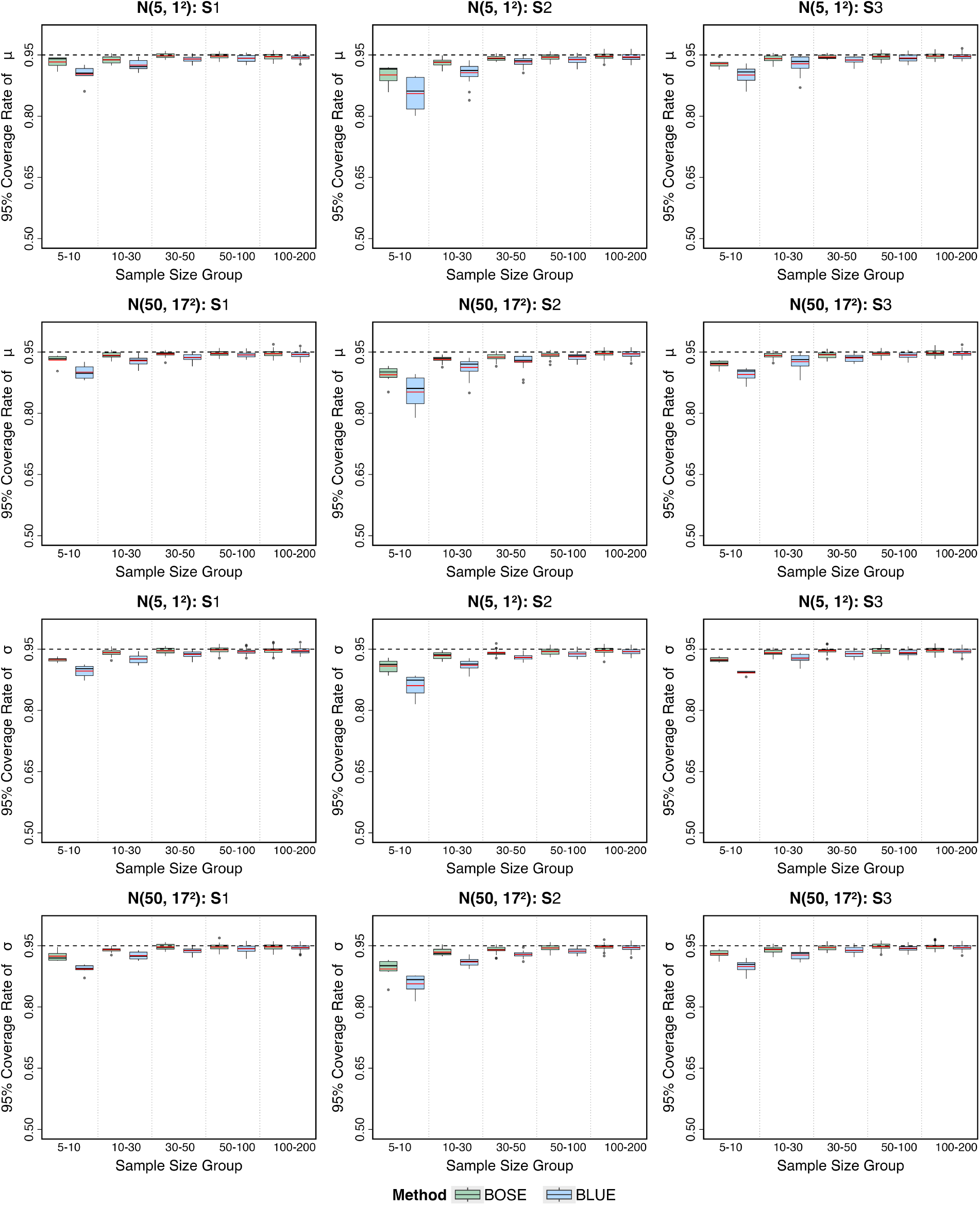
Empirical coverage rates of nominal 95% intervals for *µ* and *σ* under normal distributions *N*(5, 1^2^) (top row) and N(50, 17^2^) (bottom row), comparing BOSE credible intervals with BLUE confidence intervals across reporting scenarios *S*_1_ −*S*_3_. Each boxplot summarizes coverage rates across the evaluated sample sizes from 5 to 200, with coverage calculated over simulation replicates at each sample size. The horizontal dashed line marks the nominal 95% level, and the red lines within the boxplots indicate mean coverage rates. Coverage closer to 95% indicates better calibration.

Traditional point estimation methods (LW, BC, QE, and MLN) provide no capacity for interval estimation, leaving estimation uncertainty unquantified. BOSE addresses this limitation by deriving credible intervals from the full posterior distribution. This approach accommodates parameter constraints (e.g., *σ* > 0) and quantifies uncertainty without relying on large-sample asymptotic assumptions, providing a more robust and informative foundation for downstream meta-analytic synthesis.

We additionally examined whether the choice of quantile type affects the empirical coverage of BOSE’s credible intervals, shown in Supplementary Figure S6. Coverage remains close to the nominal 95% level across all nine quantile types for both *µ* and *σ*, with median coverage consistently falling within 0.94–0.95 regardless of the type used. Larger deviations from the nominal level were limited to a few cases in Scenario *S*_2_ for Type 3 and Type 7, but this does not correspond to any systematic shift in the median coverage for these types. Together with the RMSE results in Figure 3, this suggests that BOSE’s performance, including its interval estimation properties, is quite robust to the choice of sample quantile definition.

### 3.3 Sensitivity Analysis under Non-Normality

Figures 5 and 6 summarize mean and SD estimation, respectively, under the primary simulation settings with mild to moderate deviations from normality described in Section 3.1. Results under more extreme skewness and heavier tails are presented separately in Supplementary Figures S7 and S8. Supplementary Figures S9 and S10 report empirical coverage for interval estimation of the mean and SD under the primary settings described above. Computational failures and the RMSE threshold-based exclusions are handled uniformly across all methods, as described in Section 3.1, and the resulting NA proportions are annotated in each panel. Because averaged RMSE is computed over non-NA replications only, methods with higher NA rates may appear more favorable than their actual performance warrants. All results are averaged across sample sizes from 5 to 50.

**Figure 5:**
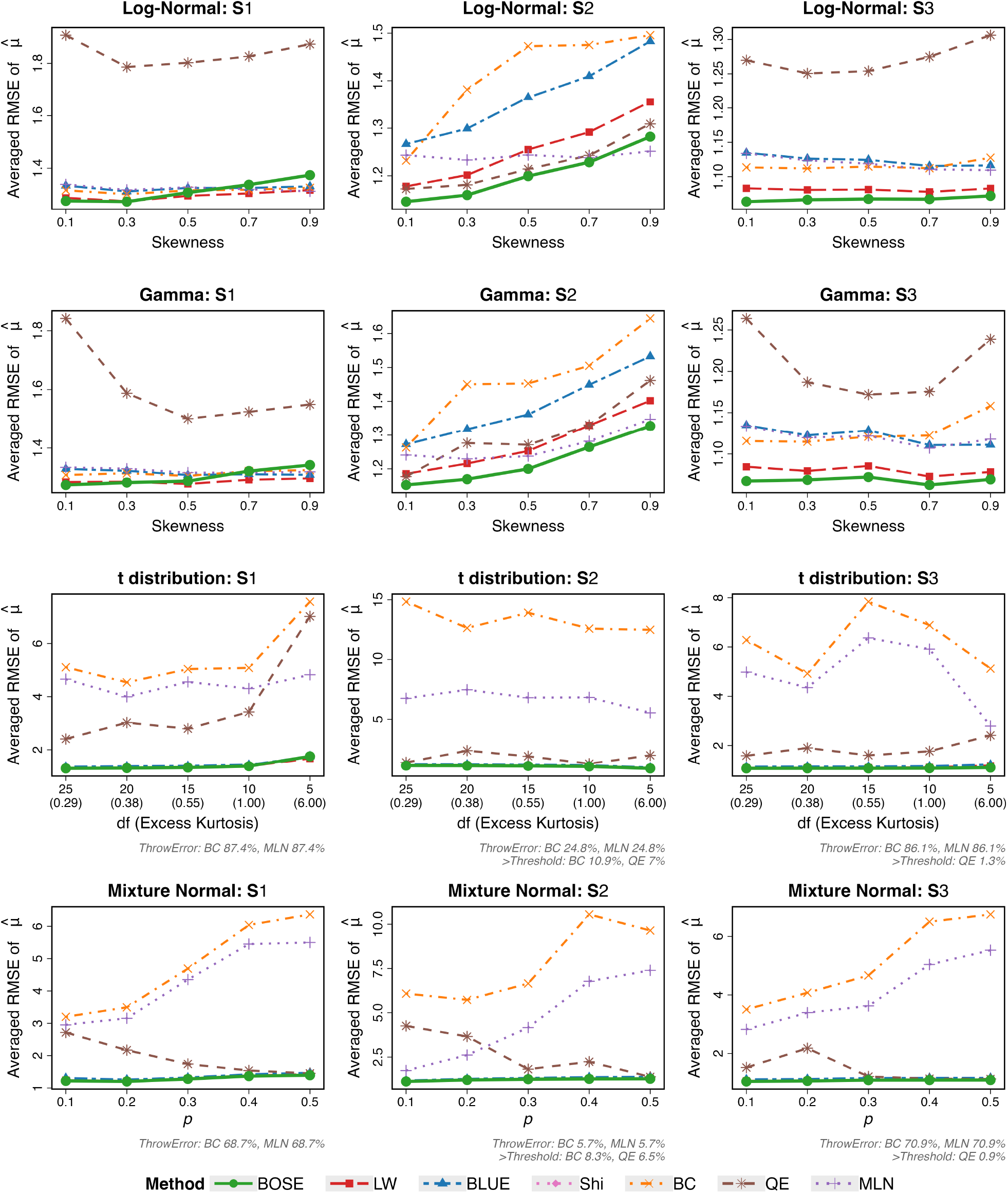
Sensitivity analysis based on averaged RMSE for mean estimation using data generated from four non-normal distributions across sample size 5 to 50: Log-Normal (1st row), Gamma (2nd row), Student’s t (3rd row), and normal mixture with two modes (4th row). The performance of BOSE (solid green line) is compared with the existing methods when applicable across varying distribution parameters (skewness, degrees of freedom/excess kurtosis, and mixing proportion *p*).

**Figure 6:**
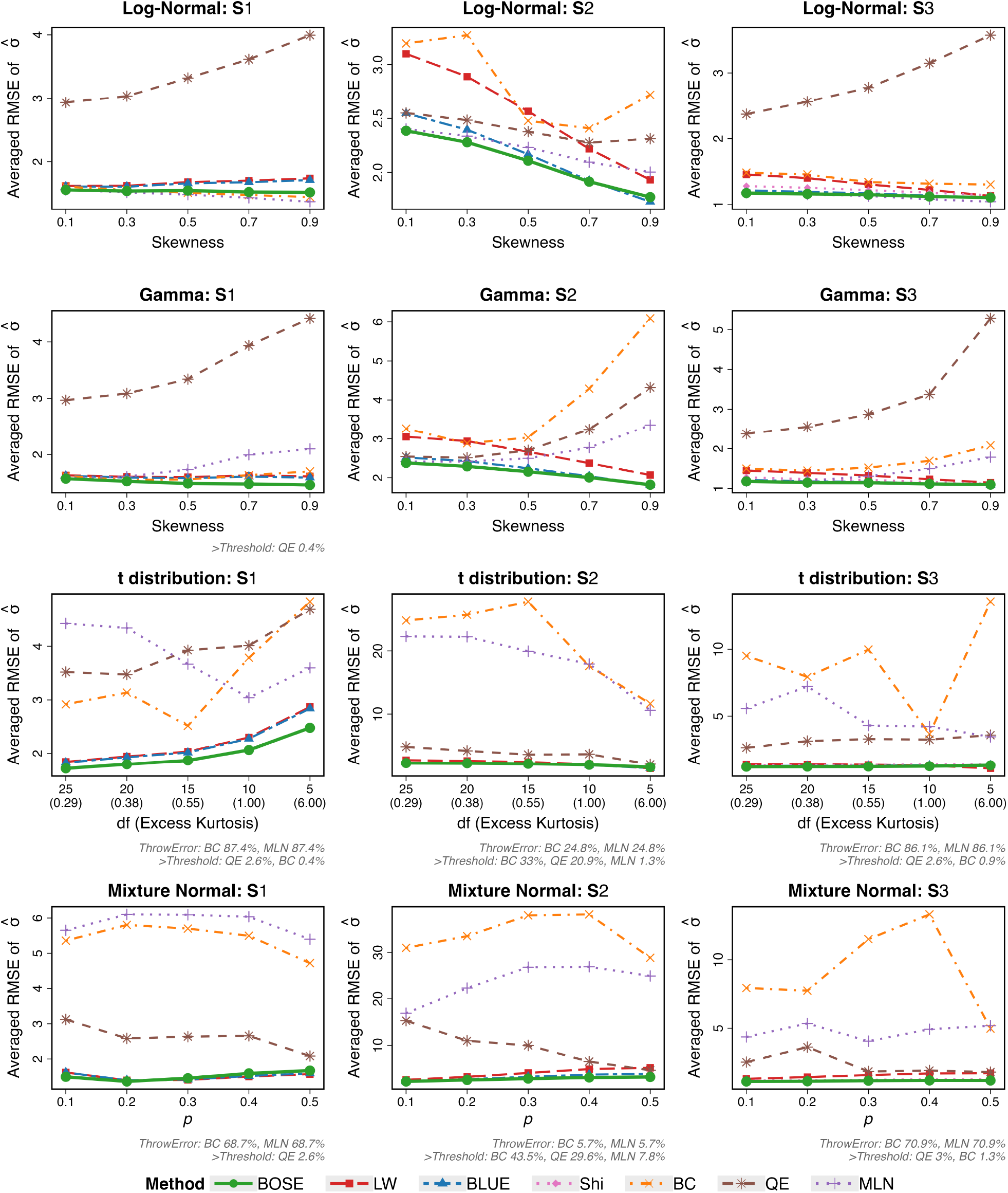
Sensitivity analysis based on averaged RMSE for SD estimation using data generated from four non-normal distributions across sample size 5 to 50: Log-Normal (1st row), Gamma (2nd row), Student’s t (3rd row), and normal mixture with two modes (4th row). The performance of BOSE (solid green line) is compared with the existing methods when applicable across varying distribution parameters (skewness, degrees of freedom/excess kurtosis, and mixing proportion *p*).

For mean estimation, Figure 5 shows that BOSE achieves the lowest or near-lowest averaged RMSE across all distributions, scenarios, and parameter settings included in the primary assessment. Among the three types of non-normal distributions, skewness exerts the largest effect: under both log-normal and gamma distributions, the RMSE of all methods rises with skewness, most steeply in *S*_2_. In *S*_1_, LW produces slightly lower RMSE than BOSE beyond skewness 0.3, though the difference remains within 0.05. Under heavy-tailed (Student’s t) and bimodal (normal mixture) distributions, normality-based methods remain clustered at low RMSE values while skewness-based methods perform considerably worse. The high NA rates of BC and MLN (exceeding 70% in most t and normal mixture panels) indicate frequent negative inputs that violate their positivity requirement. QE handles the same inputs by defaulting to a normal fit, so its apparent advantage over BC and MLN in these settings largely reflects estimation under the normal assumption.

For SD estimation, Figure 6 shows that the differences in averaged RMSE across methods are more evident. Under skewed distributions, BOSE maintains the lowest RMSE across most scenarios. For log-normal *S*_1_ and *S*_3_, MLN achieves competitive RMSE, and BC also performs well in *S*_1_; however, neither advantage carries over to the gamma distributions. A notable pattern appears under the gamma distributions: the RMSE of normality-based methods decreases as skewness grows, while skewness-based methods show the opposite trend. Under heavy-tailed (t) and bimodal (normal mixture) distributions, the limitations of skewness-based methods extend beyond their inability to handle negative inputs. In *S*_2_, where the proportion of negative summary statistics is lowest among the three scenarios (e.g., 5.7% for normal mixture), BC and MLN still produce RMSE values above 10, much worse than in *S*_1_ and *S*_3_ where their NA rates exceed 65%. This indicates that BC and MLN are not only constrained by the positivity requirement but also sensitive to the distributional shape itself: even when they can produce estimates, those estimates are unreliable under the tested heavy-tailed and bimodal settings.

In the primary simulation settings with mild to moderate deviations from normality, BOSE performed comparably to or better than the other methods in estimating both the mean and SD. We next examined performance under more extreme skewness and heavier tails, with results presented separately for mean estimation (Supplementary Figure S7) and SD estimation (Supplementary Figure S8). Under more extreme skewness, the skewness-based method MLN achieved the lowest RMSE for mean estimation among all methods evaluated. BOSE’s RMSE for mean estimation increased more rapidly with skewness than did those of the other two normality-based methods, LW and BLUE, indicating a limitation of BOSE in highly skewed settings. BOSE remained competitive for SD estimation under extreme skewness and for both mean and SD estimation under heavier-tailed distributions, with RMSEs only slightly higher than those of LW and BLUE. Consistent with our expectations, these findings suggest that MLN is preferable to BOSE for mean estimation under the highly skewed distributions examined here.

Taken together, BOSE delivers the best or near-best averaged RMSE for both 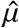 and 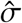 across most tested non-normal settings, with the exception of mean estimation under more extreme skewness *κ* ∈ {1, 2, 3}, where skewness-based method MLN is preferable. The advantage over competitors is more pronounced for SD estimation than for mean estimation, as competing methods degrade more rapidly for SD estimation while BOSE remains stable.

For the same primary simulation settings shown in Figures 5 and 6, we also evaluated the empirical coverage of BOSE’s 95% credible intervals for *µ* and *σ* under the four non-normal distribution families, alongside BLUE’s corresponding confidence intervals (Supplementary Figures S9 and S10).

For *µ*, BOSE generally achieves coverage comparable to or closer to 95% than BLUE. The improvement is most apparent under the log-normal and gamma distributions in Scenario *S*_2_, although both methods show increasing under-coverage as skewness increases. Under Student’s t distributions, coverage for *µ* remains relatively stable, with BOSE closer to the nominal level in Scenario *S*_2_. Under normal mixtures, the methods perform similarly overall, with coverage depending on the mixture proportion and reporting scenario.

For *σ*, BOSE also generally provides comparable or improved coverage, particularly in Scenario *S*_2_. Coverage declines with increasing skewness, and, more markedly, with heavier tails. At five degrees of freedom, both methods exhibit substantial under-coverage despite BOSE’s improvement over BLUE in Scenario *S*_2_. Under normal mixtures, coverage for both methods generally approaches 95% as the mixing proportion increases in Scenarios *S*_2_ and *S*_3_.

Overall, within the primary simulation settings, BOSE often provides comparable or better empirical calibration than BLUE, but neither method consistently maintains nominal coverage under non-normality. The under-coverage observed for *σ* under heavy-tailed distributions highlights a limitation of interval estimation based on the assumed normal model.

## 4 Real Data Analysis

### 4.1 Serum Vitamin D Levels and Tuberculosis Risk

We demonstrate the practical utility of BOSE using data from a systematic review examining the association between low serum vitamin D levels and active tuberculosis (TB) risk, following the work of Luo et al.^5^ and Yang, Hutson, and Wang^7^. The dataset comprises five independent studies extracted from the systematic review by Nnoaham and Clarke^18^, of which three (Studies 1–3) reported only sample medians and ranges (Scenario *S*_1_) and two (Studies 4–5) provided means and SDs directly. A detailed summary of study characteristics is reproduced in Supplementary Table S3 for convenience, and estimation results across all methods are provided in Table 1. The mean difference (MD) and standardized mean difference (SMD) were used to compare serum vitamin D levels between the treatment and control groups.

**Table 1:**
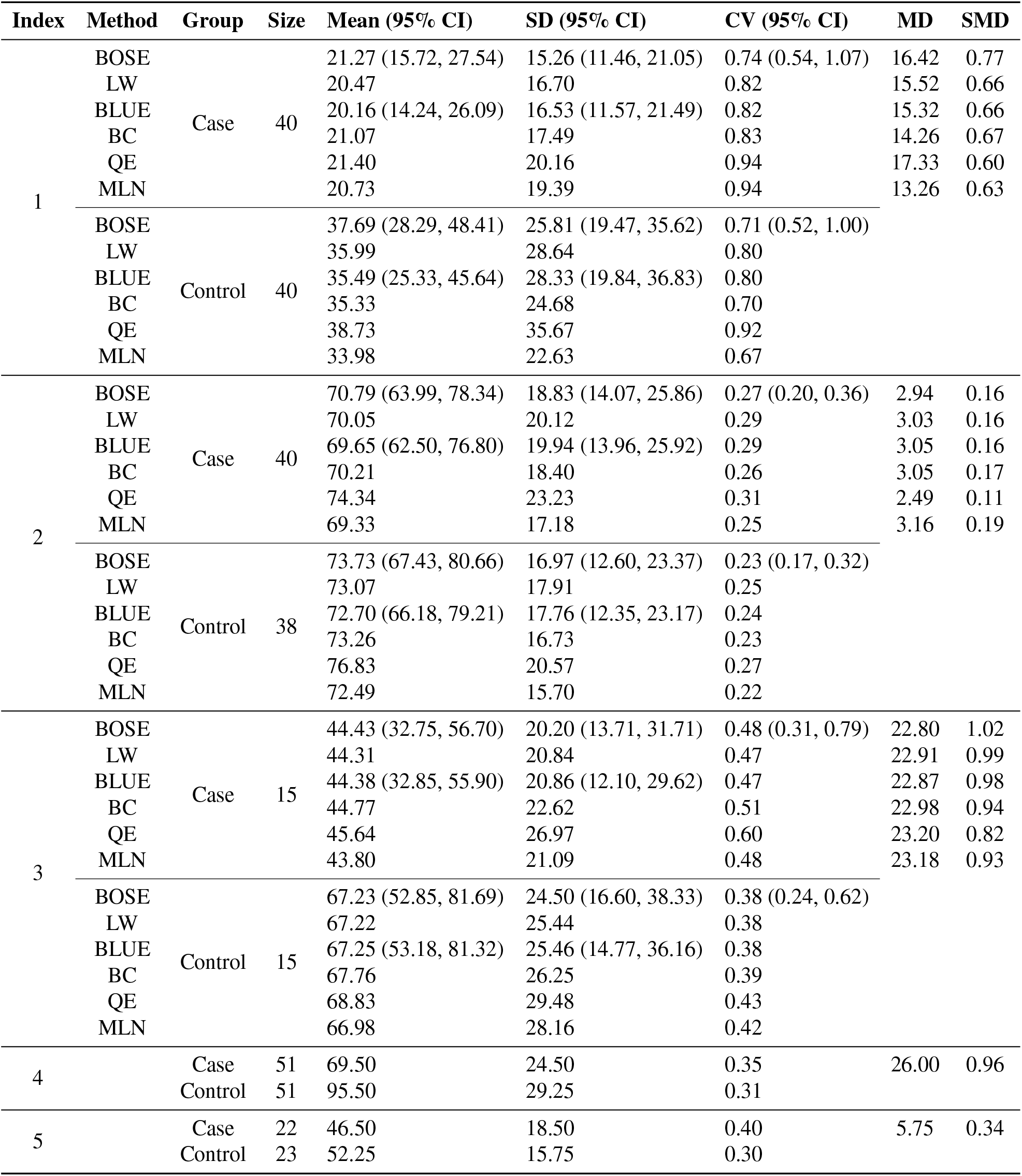
Data Example I: estimated effect sizes of low serum vitamin D in tuberculosis. MD stands for mean difference and SMD stands for standardized mean difference between case and control groups.

For Studies 1–3, where estimation from summary statistics is required, point estimates across methods show some variation. For example, mean estimates for Study 1 Case range from 20.16 (BLUE) to 21.40 (QE), and SD estimates range from 15.26 (BOSE) to 20.16 (QE). These differences can propagate into MD and SMD calculations and may affect the conclusions of the subsequent meta-analysis.

BOSE provides empirically calibrated credible intervals for all estimated quantities directly from the posterior distribution. Traditional point estimation methods (e.g., LW, BC, QE, and MLN) do not provide this capability, leaving estimation uncertainty unquantified. BLUE provides confidence intervals via asymptotic approximations 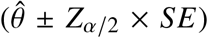, which assume normality and yield symmetric intervals; this construction may be less suitable in small samples for inherently non-negative, skewed parameters such as *σ*. In contrast, BOSE derives intervals directly from the empirical posterior, which can accommodate asymmetry when present. For Study 3 (*n* = 15), the BOSE SD interval for the Case group is (13.71, 31.71), compared with BLUE’s (12.10, 29.62) centered at its SD estimate 20.86. The two intervals are of comparable width, but BOSE’s interval is asymmetric, consistent with the right-skewed marginal posterior of the variance parameter.

The contour plots in Figure 7 visualize this uncertainty structure. For the first two studies (*n* = 38 to 40), the joint posterior distributions are tightly concentrated, with the BOSE point estimates (red cross) located within the highest-density regions. For Study 3, with a smaller sample size (*n* = 15), the contours are more diffuse, reflecting the greater estimation uncertainty typically associated with smaller samples. Across the panels, the frequentist LW estimates (blue triangles), which are used to initialize the BOSE grid, exhibit some deviation from the final posterior highest-density regions, particularly along the vertical *σ*-axis, illustrating how the posterior update can shift the estimate away from its initial value. The tilted, elliptical shape of the contours reflects the correlation between *µ* and *σ*, and the asymmetric spacing of the credible interval bounds around the point estimates reflects the right-skewed marginal posterior of the variance parameter.

**Figure 7:**
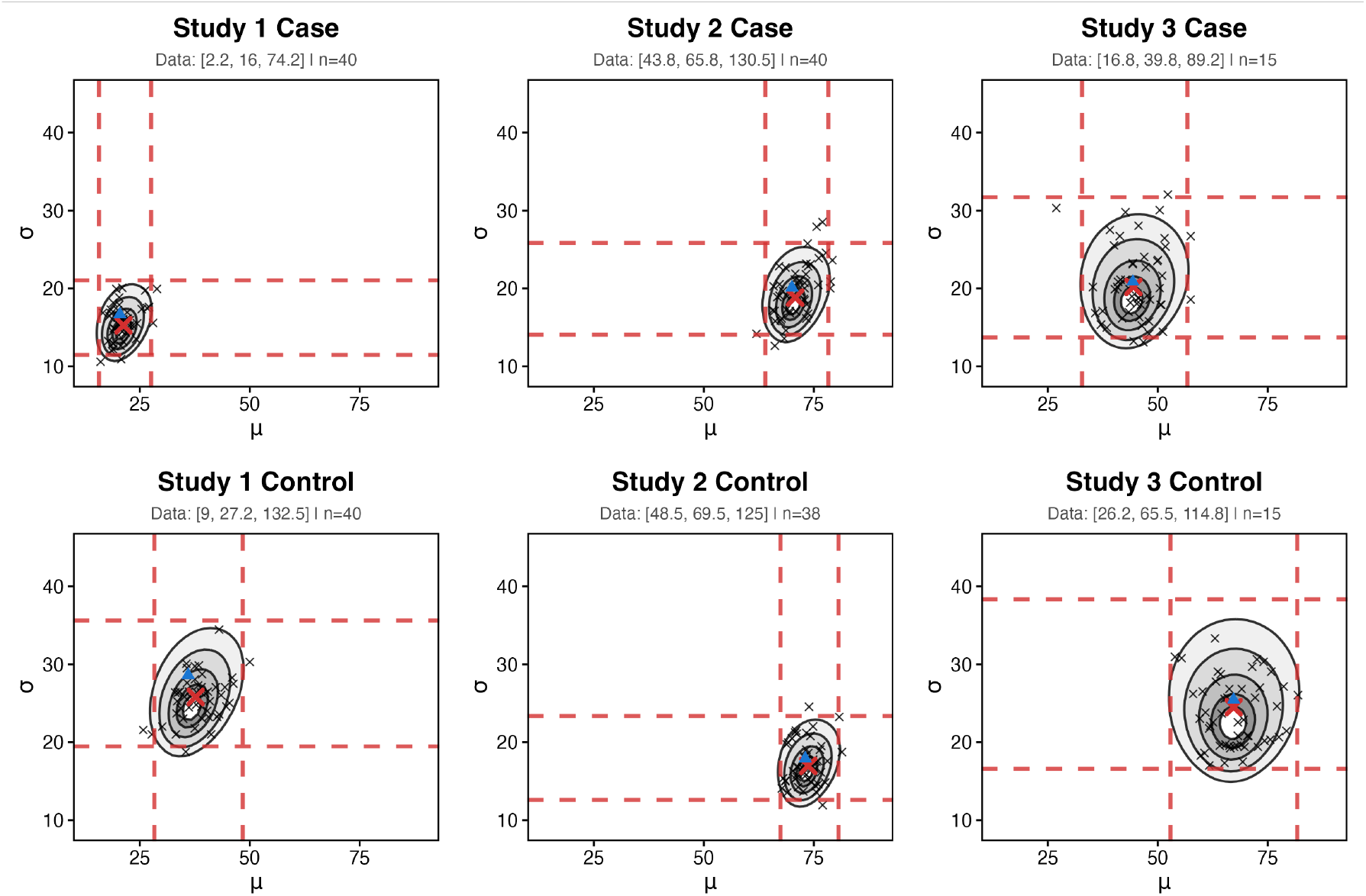
Data Example I: contour plots of the joint posterior distributions of (*µ, σ*) for studies 1–3 across Case (top row) and Control (bottom row) groups. Gray filled contours with overlaid lines represent the posterior density, with 50 random draws (black crosses) shown out of 1000 total posterior samples. The red cross denotes the marginal posterior point estimates (posterior mean for *µ* and posterior median for *σ*) derived from the proposed BOSE method. The blue triangle indicates the initial estimates obtained from the LW method. The red dashed lines mark the marginal 95% credible intervals for *µ* and *σ*.

A further advantage of BOSE is its ability to quantify uncertainty in non-linear derived quantities, such as the coefficient of variation (*CV* = *σ*/*µ*), obtained via Monte Carlo sampling from the joint posterior of (*µ, σ*). For Study 1 Case, where the CV estimate is 0.74, BOSE provides a 95% credible interval of (0.54, 1.07), a range wide enough to include values exceeding 1, indicating that the relative variability of serum Vitamin D in this group may exceed the mean itself. This degree of uncertainty would not be apparent from point estimates alone. By contrast, Study 2 Case yields a CV of 0.27 with a narrower interval of (0.20, 0.36), indicating greater precision in the CV estimate. The contrast between these two studies, both with *n* = 40, illustrates that sample size alone does not determine estimation precision; the underlying data structure also plays a role. The non-overlapping CV intervals between Study 1 Case (0.54, 1.07) and Study 2 Case (0.20, 0.36) are suggestive of heterogeneity in relative variability across studies, a distinction that point estimates alone would not reveal.

### 4.2 COVID-19 and Liver Dysfunction

Our second example uses data from a meta-analysis by Wu and Yang^19^ examining the impact of COVID-19 on liver dysfunction, as measured by alanine aminotransferase (ALT) levels, following the work of Shi et al.^17^. The dataset comprises four studies, all reporting summary statistics under Scenario *S*_2_. A summary of study characteristics is reproduced in Supplementary Table S4 for convenience. Forest plots comparing three different analytical approaches as detailed below are displayed in Figure 8, presenting point estimates for each study; the corresponding credible intervals for the mean, SD, and Coefficient of Variation (CV) obtained by BOSE are reported separately in Table 2.

**Table 2:** Data Example II: BOSE posterior point estimates and 95% credible intervals.

| Index | Study | Group | Size | Mean (95% CI) | SD (95% CI) | CV (95% CI) |
| --- | --- | --- | --- | --- | --- | --- |
| 1 | Chen et al.<sup>21</sup> | Non-Survivor | 113 | 31.16 (26.78, 35.50) | 21.63 (17.66, 27.09) | 0.69 (0.55, 0.92) |
| 2 | Du et al.<sup>22</sup> |  | 21 | 28.08 (21.56, 34.50) | 12.70 (8.23, 22.08) | 0.45 (0.29, 0.85) |
| 3 | Wang et al.<sup>20</sup> |  | 65 | 31.07 (24.81, 37.25) | 22.87 (17.62, 30.91) | 0.74 (0.55, 1.11) |
| 4 | Zhou et al.<sup>23</sup> |  | 54 | 38.38 (32.48, 44.20) | 19.66 (14.83, 27.31) | 0.51 (0.38, 0.74) |
| 1 | Chen et al.<sup>21</sup> | Survivor | 161 | 22.39 (20.26, 24.49) | 12.87 (10.84, 15.52) | 0.58 (0.48, 0.71) |
| 2 | Du et al.<sup>22</sup> |  | 158 | 25.74 (22.41, 29.06) | 19.69 (16.58, 23.76) | 0.77 (0.63, 0.97) |
| 3 | Wang et al.<sup>20</sup> |  | 274 | 29.43 (26.94, 31.88) | 19.21 (16.83, 22.15) | 0.65 (0.56, 0.78) |
| 4 | Zhou et al.<sup>23</sup> |  | 135 | 27.35 (23.96, 30.68) | 18.22 (15.15, 22.33) | 0.67 (0.54, 0.86) |

**Figure 8:**
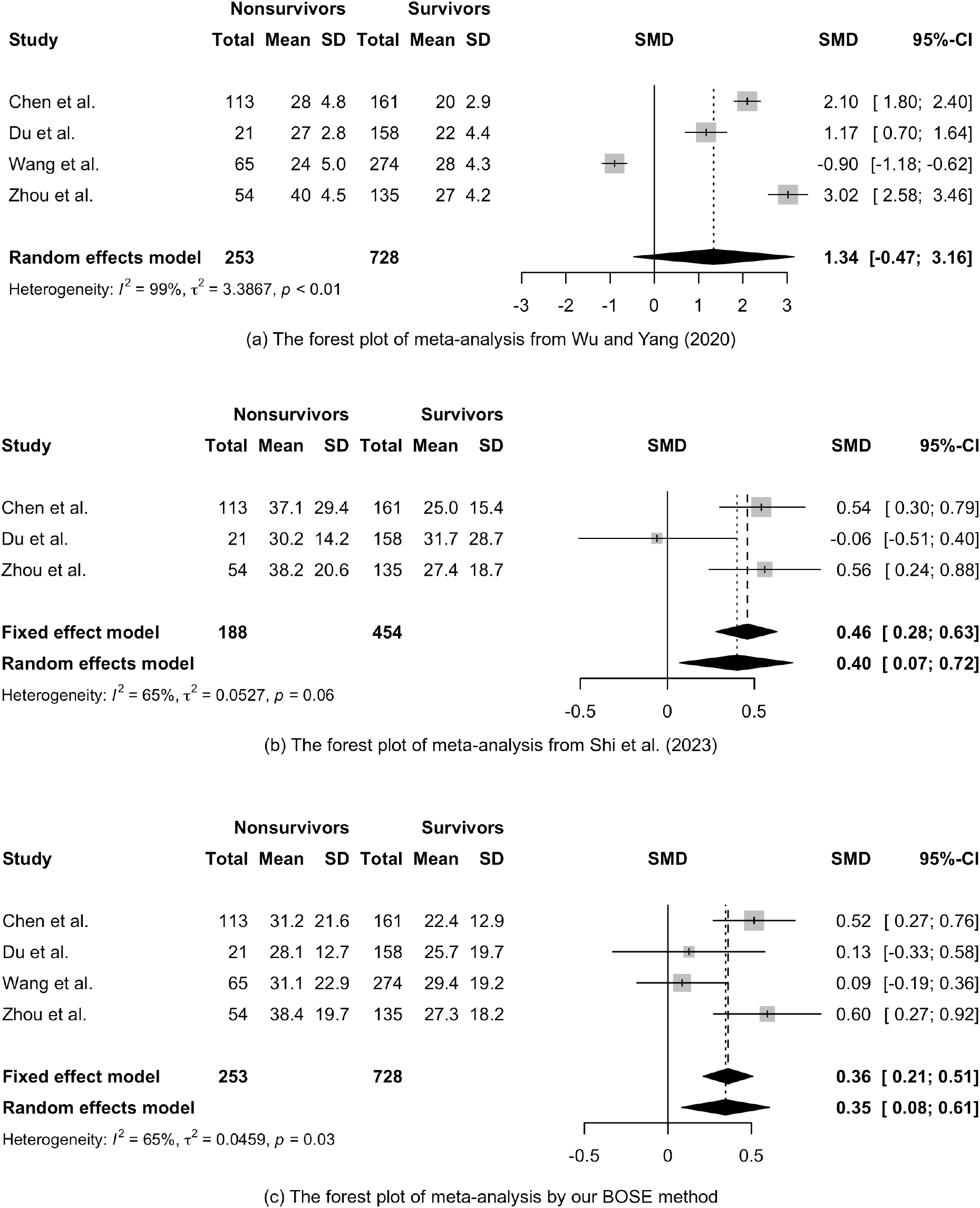
Data Example II: forest plots comparing meta-analyses of standardized mean differences (SMDs) between non-survivors and survivors using three different approaches. Panels (a) and (b) are reproduced from Shi et al.^17^, showing the original meta-analysis by Wu and Yang^19^ and a re-analysis excluding Wang et al.^20^, respectively. Panel (c) presents our results using the proposed BOSE method.

Wu and Yang^19^ applied the Hozo, Djulbegovic, and Hozo^3^ transformation to estimate the missing sample means and SDs, yielding an SMD of 1.34 (95% CI: −0.47, 3.16) and a conclusion of no significant effect under the random-effects (RE) model. However, the implausibly small SD estimates produced by this transformation (ranging from 2.8 to 5.0) artificially inflate individual SMDs and drive an extreme heterogeneity index of *I*^2^ = 99%, limiting the reliability of the synthesis. Shi et al.^17^ subsequently addressed this by introducing a skewness testing procedure. Their analysis found that the non-survivor group in Wang et al.^20^ remained significantly skewed even after a log-scale transformation, leading to the study’s justifiable exclusion. Synthesizing the remaining three studies yielded an SMD of 0.40 (95% CI: 0.07, 0.72) and reduced the heterogeneity index to *I*^2^ = 65% under the RE model. While this approach provides a statistically significant result, it excludes 339 participants based on distributional assumptions rather than data quality.

Applying BOSE directly to all four studies without any preliminary testing yields an SMD of 0.35 (95% CI: 0.08, 0.61) with the same heterogeneity (*I*^2^ = 65%) as in the second approach. Furthermore, the posterior credible intervals provide quantitative context for the exclusion decision made by Shi et al.^17^. As summarized in Table 2, the CV for the Wang et al.^20^ non-survivor group is 0.74, with a 95% credible interval of (0.55, 1.11). The upper bound exceeding 1 suggests substantial within-group variability, consistent with this group’s exclusion under the log-scale normality test applied by Shi et al.^17^.

A closer examination of individual study results highlights the instability of parameter recovery in settings with highly imbalanced or small-arm sample sizes. For Du et al.^22^, although the survivors group is sufficiently large (*n* = 158), the non-survivor group comprises only *n* = 21 participants. In such imbalanced scenarios, the substantial estimation uncertainty within the smaller arm dominates the overall comparative metric. Consequently, the estimated means and SDs for this smaller group fluctuate drastically across methodologies, causing the resulting SMD estimates to diverge widely: 1.17 under Wu and Yang^19^, −0.06 under Shi et al.^17^, and 0.13 under BOSE. The negative SMD from the Shi et al.^17^ estimator is driven by an inversion of the estimated group means (30.2 for non-survivors versus 31.7 for survivors). This reversal implies that survivors had higher ALT levels than non-survivors, a direction inconsistent with both other approaches and with the overall clinical expectation regarding liver dysfunction severity. This phenomenon illustrates how differing estimation frameworks can yield unstable and potentially misleading point estimates when applied to imbalanced data.

Finally, this example illustrates that the choice of estimation methods can lead to qualitatively different clinical interpretations. BOSE’s credible intervals offer additional diagnostic information beyond the point estimates: as shown in Table 2, credible interval widths vary across studies, with Chen et al.^21^ showing comparatively narrow intervals across both the non-survivor and survivor subgroups, while Du et al.^22^, Wang et al.^20^, and Zhou et al.^23^ show wider intervals in both subgroups; the latter signal greater estimation uncertainty and warrant more caution when interpreting their point estimates.

## 5 Conclusions and Discussion

We introduced BOSE, a novel Bayesian method designed to estimate sample means and SDs from heterogeneous summary statistics in meta-analyses. Our simulations and real-world data applications showed that BOSE achieves competitive or improved performance relative to existing methods across a broad range of settings, while most competing approaches provide only point estimates without quantifying uncertainty.

Under the normality assumption, BOSE achieves lower averaged RMSE for both mean and SD estimation, with the advantage more pronounced for SD *σ* and in small-sample settings. Beyond point estimation, BOSE provides well-calibrated credible intervals at multiple nominal levels, derived directly from the posterior distribution rather than asymptotic approximations, giving researchers a more reliable measure of estimation uncertainty prior to committing estimates to downstream meta-analytic calculations.

BOSE further demonstrates robustness against departures from normality. Across skewed, heavy-tailed, and bimodal distributions, it maintains stable and competitive performance while many competing methods suffer from increasing instability or outright numerical failure. Existing skewness-based methods in particular are fundamentally inapplicable when quantile values are negative, rendering them unreliable in precisely the settings for which they were designed.

The real-world applications highlight a broader implication for evidence synthesis: estimates reconstructed from limited summary statistics benefit from accompanying measures of uncertainty. As shown in both the tuberculosis and COVID-19 examples, BOSE’s posterior credible intervals complement point estimates and help researchers identify study-level estimates that warrant closer scrutiny before use in downstream meta-analysis.

These results suggest that BOSE is a practical default choice for meta-analysis under normality and mild-to-moderate departures from normality. Applicable directly to any combination of reported order statistics without prior distributional knowledge or formal testing procedures, it offers a robust and accessible option for evidence synthesis when the distributional properties of primary studies are unknown or uncertain. To facilitate the broad accessibility and practical application of our method, we have developed an interactive web-based tool (available at https://bose-project.shinyapps.io/shinyapp/). This tool computes point estimates alongside their corresponding credible intervals and recommends estimation strategies, making uncertainty quantification accessible to practitioners regardless of their statistical background.

The current implementation focuses on study-level estimation, treating each study independently. A natural and important extension is the development of a fully integrated Bayesian hierarchical model that jointly estimates study-level parameters and between-study heterogeneity within a unified framework, allowing uncertainty from the estimation step to propagate directly into the meta-analytic pooling step. This represents the primary direction of our ongoing work.

## Code Availability

An interactive web-based tool is available at https://bose-project.shinyapps.io/shinyapp/. The R code used in this study is available at https://github.com/wenqisi-pan/BOSE-bayesian-order-statistics-estimator.

