## Supplementary Material for "BOSE: A Bayesian Order Statistics-Based Estimator for Recovering the Sample Mean and Standard Deviation from the Median, Range, and/or Interquartile Range"

### 1 Implementation Details

Algorithm 1 details our adaptive grid strategy for Scenario $S_3$ (the five-number summary). This algorithm first constructs a coarse grid to locate the high-density Region of Interest (ROI) and subsequently deploys a fine grid strictly within this bounding box to achieve high-precision numerical integration.

**Algorithm 1** BOSE with Adaptive Grid for Scenario $S_3$

**Require:** Input data $\mathcal{S}_3 = \{a, q_1, m, q_3, b; n\}$

**Step 1:** Obtain initial estimates for $\mu$ and $\sigma$ using LW method

$\hat{\mu}_{init} = \hat{\mu}_0 = \left(\frac{2.2}{2.2+n^{0.75}}\right)\frac{a+b}{2} + \left(0.7 - \frac{0.72}{n^{0.55}}\right)\frac{q_1+q_3}{2} + \left(0.3 + \frac{0.72}{n^{0.55}} - \frac{2.2}{2.2+n^{0.75}}\right)m$

$\hat{\sigma}_{init} = \hat{\sigma}_0 = \frac{b-a}{4\Phi^{-1}\left(\frac{n-0.375}{n+0.25}\right)} + \frac{q_3-q_1}{4\Phi^{-1}\left(\frac{0.75n-0.125}{n+0.25}\right)}$

**Step 2:** Define data-centered initial search region

Let $\mu_L = \hat{\mu}_0 - z\hat{\sigma}_0$, $\mu_R = \hat{\mu}_0 + z\hat{\sigma}_0$; $\sigma_L = \hat{\sigma}_0/z$, $\sigma_R = z\hat{\sigma}_0$

**Step 3 (Stage 1):** Construct coarse grid $G_c \times G_c$ to locate high-density region

Set $\mathcal{G}_\mu^{(c)} = \{\mu_L + \frac{i-1}{G_c-1}(\mu_R - \mu_L)\}_{i=1}^{G_c}$; $\mathcal{G}_\sigma^{(c)} = \{\sigma_L + \frac{j-1}{G_c-1}(\sigma_R - \sigma_L)\}_{j=1}^{G_c}$

Compute normalized posterior weights $\mathcal{W}_{ij}^{(c)} \propto p(\mu_i^{(c)}, \sigma_j^{(c)} | \mathcal{S}_3)$ for $i, j = 1, \ldots, G_c$

**Step 4:** Identify Region of Interest (ROI) containing 99% of posterior mass

Sort weights in descending order: $\mathcal{W}_{(1)}^{(c)} \geq \mathcal{W}_{(2)}^{(c)} \geq \cdots \geq \mathcal{W}_{(G_c^2)}^{(c)}$

Find smallest $K$ such that $\sum_{k=1}^{K} \mathcal{W}_{(k)}^{(c)} \geq 0.99$; Let $\mathcal{W}_{\text{thresh}} = \mathcal{W}_{(K)}^{(c)}$ be the threshold weight

Define high-density region: $\mathcal{R} = \{(i, j) : \mathcal{W}_{ij}^{(c)} \geq \mathcal{W}_{\text{thresh}}\}$

Extract ROI bounds $[\mu_{\min}^{ROI}, \mu_{\max}^{ROI}] \times [\sigma_{\min}^{ROI}, \sigma_{\max}^{ROI}]$ from high-density grid points:

$\mu_{\min}^{\text{ROI}} = \min\{\mu_i^{(c)} : (i, j) \in \mathcal{R}\}$, $\mu_{\max}^{\text{ROI}} = \max\{\mu_i^{(c)} : (i, j) \in \mathcal{R}\}$

$\sigma_{\min}^{\text{ROI}} = \min\{\sigma_j^{(c)} : (i, j) \in \mathcal{R}\}$, $\sigma_{\max}^{\text{ROI}} = \max\{\sigma_j^{(c)} : (i, j) \in \mathcal{R}\}$

Expand ROI by 10%:

$\mu_L^{(f)} = \mu_{\min}^{\text{ROI}} - 0.1(\mu_{\max}^{\text{ROI}} - \mu_{\min}^{\text{ROI}})$, $\mu_R^{(f)} = \mu_{\max}^{\text{ROI}} + 0.1(\mu_{\max}^{\text{ROI}} - \mu_{\min}^{\text{ROI}})$

$\sigma_L^{(f)} = \sigma_{\min}^{\text{ROI}} - 0.1(\sigma_{\max}^{\text{ROI}} - \sigma_{\min}^{\text{ROI}})$, $\sigma_R^{(f)} = \sigma_{\max}^{\text{ROI}} + 0.1(\sigma_{\max}^{\text{ROI}} - \sigma_{\min}^{\text{ROI}})$

**Step 5 (Stage 2):** Construct fine grid $G_f \times G_f$ within ROI

Set $\mathcal{G}_\mu^{(f)} = \{\mu_L^{(f)} + \frac{i-1}{G_f-1}(\mu_R^{(f)} - \mu_L^{(f)})\}_{i=1}^{G_f}$; $\mathcal{G}_\sigma^{(f)} = \{\sigma_L^{(f)} + \frac{j-1}{G_f-1}(\sigma_R^{(f)} - \sigma_L^{(f)})\}_{j=1}^{G_f}$

Compute normalized posterior weights $\mathcal{W}_{ij}^{(f)} \propto p(\mu_i^{(f)}, \sigma_j^{(f)} | \mathcal{S}_3)$ for $i, j = 1, \ldots, G_f$

**Step 6:** Compute marginal posterior distributions

$p(\mu_i^{(f)} \mid \mathcal{S}_3) = \sum_{j=1}^{G_f} \mathcal{W}_{ij}^{(f)}$ for $i = 1, \ldots, G_f$

$p(\sigma_j^{(f)} \mid \mathcal{S}_3) = \sum_{i=1}^{G_f} \mathcal{W}_{ij}^{(f)}$ for $j = 1, \ldots, G_f$

**Step 7:** Compute BOSE estimates via direct numerical integration

*Point estimates:*

$\hat{\mu}_{\text{BOSE}} = \sum_{i=1}^{G_f} \mu_i^{(f)} \cdot p(\mu_i^{(f)} \mid \mathcal{S}_3)$ *(posterior mean)*

$\hat{\sigma}_{\text{BOSE}} = F_\sigma^{-1}(0.5)$ *(posterior median via CDF inversion)*

*Credible intervals via CDF inversion:*

$\text{CI}_\mu^{95\%} = \left[F_\mu^{-1}(0.025),\ F_\mu^{-1}(0.975)\right]$

$\text{CI}_\sigma^{95\%} = \left[F_\sigma^{-1}(0.025),\ F_\sigma^{-1}(0.975)\right]$

where $F_\mu(x) = \sum_{\mu_i^{(f)} \leq x} p(\mu_i^{(f)} \mid \mathcal{S}_3)$ and $F_\sigma(x) = \sum_{\sigma_j^{(f)} \leq x} p(\sigma_j^{(f)} \mid \mathcal{S}_3)$

and $F^{-1}(\alpha)$ denotes the quantile computed via linear interpolation on the grid

**Ensure:** Point estimates and credible intervals: $\mathbf{\Theta}(\mathcal{S}_3) = \{\hat{\mu}_{\text{BOSE}}, \hat{\sigma}_{\text{BOSE}}, \text{CI}_\mu^{95\%}, \text{CI}_\sigma^{95\%}\}$

#### 2 Sensitivity Analysis of Hyperparameters

To evaluate the robustness of the BOSE framework against weakly informative prior choices, we conducted a sensitivity analysis under the normality assumption. This analysis systematically perturbed the hyperparameters controlling the prior distributions of $\mu$ and $\sigma$. The results for various combinations of the uniform prior range $z \in \{3, 5, 10\}$ and the inverse-gamma shape/scale parameters $\alpha = \beta \in \{0.001, 0.01, 0.1\}$ are presented in Figure S1. Across all tested scenarios and sample sizes, the averaged Relative Mean Squared Errors (RMSEs) for both the mean and standard deviation (SD) estimation remained stable. The highly overlapping performance trajectories confirm that BOSE's performance is robust to variations in the weakly informative prior specifications.

In addition to the prior hyperparameters, we examined the sensitivity of BOSE to the grid resolution used in the two-stage adaptive grid approximation. Figure S2 presents the averaged RMSE across four grid-size configurations, $(G_c, G_f) \in \{(64, 128), (125, 250), (200, 400), (256, 512)\}$, and three padding percentages (5%, 10%, and 20%) applied when constructing the fine grid. As with the prior sensitivity analysis, the resulting RMSE trajectories are nearly indistinguishable across all twelve combinations, confirming that BOSE's performance is robust to the specific choice of grid resolution and padding percentage within this range.

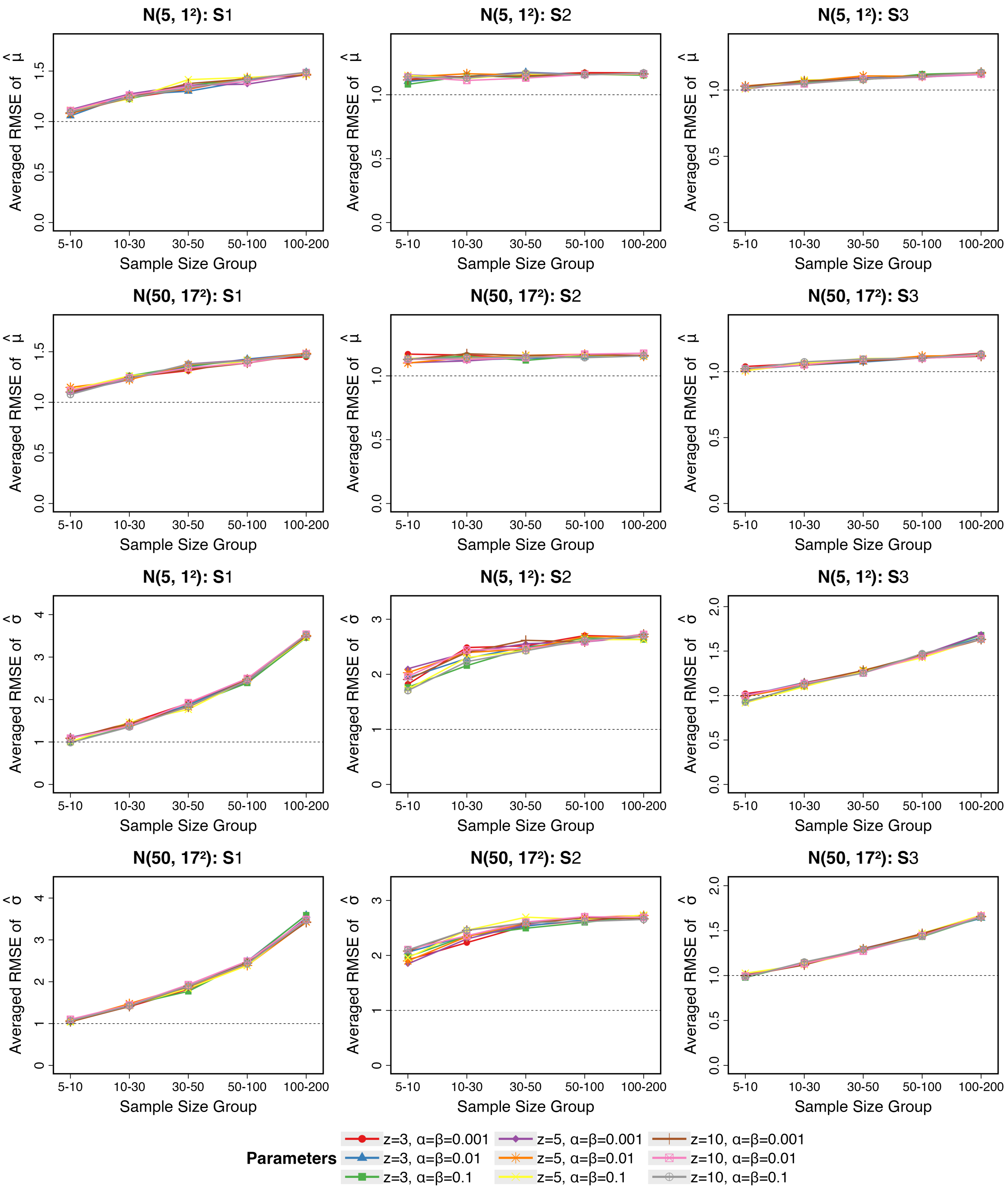


Figure S1: Sensitivity analysis of hyperparameter choices for the proposed BOSE method. Averaged RMSEs for mean estimation (top two rows) and SD estimation (bottom two rows) are evaluated across five sample size groups using data generated from normal distributions under three common data reporting scenarios $S_1 - S_3$. The analysis compares nine hyperparameter combinations: $z \in \{3, 5, 10\}$ (controlling the uniform prior range for $\mu$) and $\alpha = \beta \in \{0.001, 0.01, 0.1\}$ (shape and scale parameters for the inverse-gamma prior on $\sigma^2$).

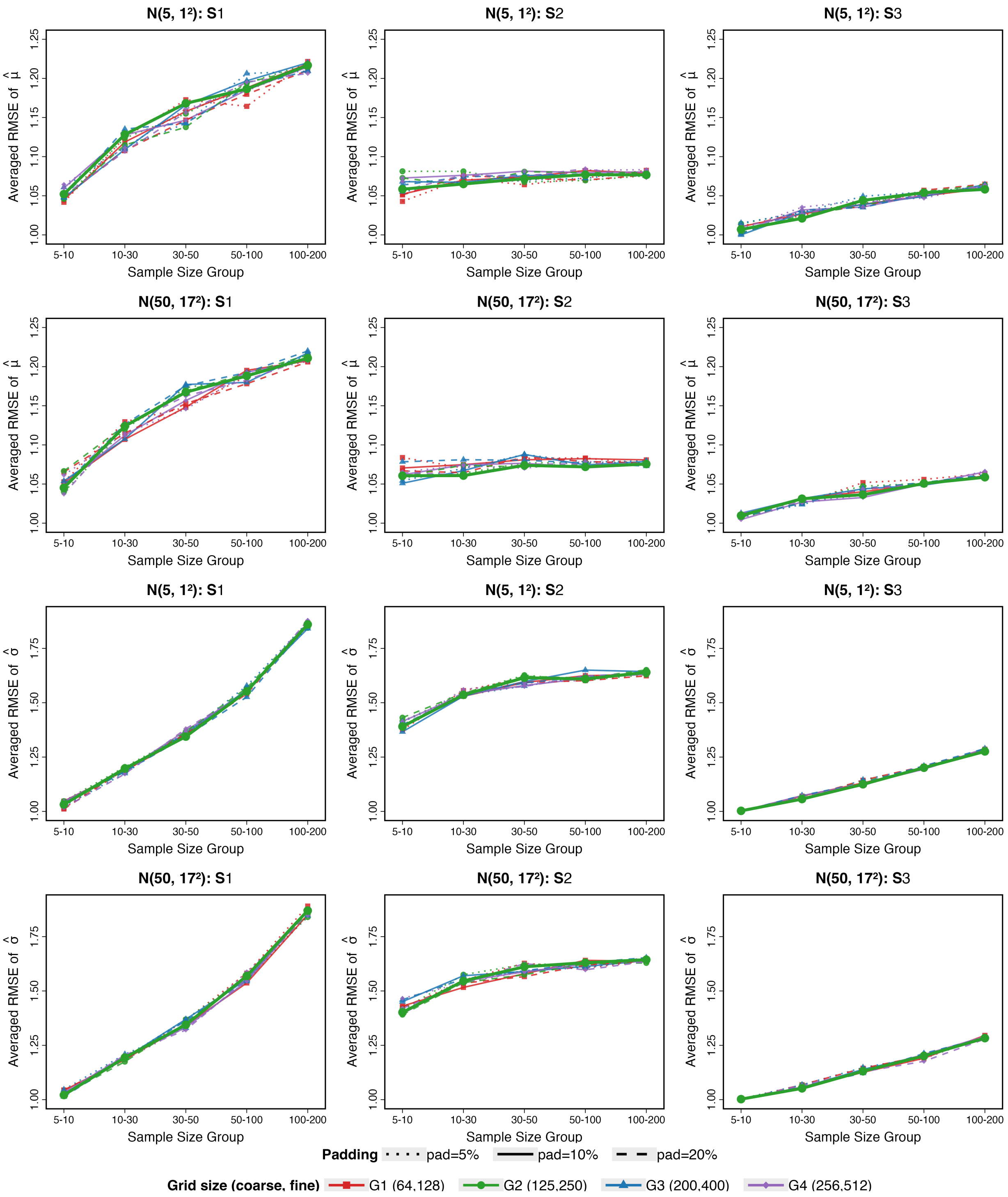


Figure S2: Sensitivity analysis of grid resolution and padding percentage based on averaged RMSE for mean estimation (top two rows) and SD estimation (bottom two rows), using data generated from normal distributions $N(5,1^2)$ and $N(50,17^2)$ under three common data reporting scenarios $S1$–$S3$. Four grid-size configurations are compared under three padding percentages applied when constructing the fine grid.

### 3 Detailed RMSE Results under Normality

Table S1 and Table S2 report the exact averaged RMSE values underlying Figure 1 of the main text, for the mean and SD, respectively. Each table is organized by sample-size group and data reporting scenario, allowing precise numerical values in addition to the visual trends shown in Figure 1.

To further confirm that the sample-size groupings used in Figures 1 and 2 do not obscure any sample-size-specific patterns, Figure S3 reports RMSE separately for each individual sample size, rather than as group averages. Simulations were run for every integer sample size from n = 5 to 200; for readability, results are displayed at every 10th value of n. The resulting trends are smooth and continuous across the full range of sample sizes, consistent with the grouped averages reported in Figures 1 and 2.

Table S1: Averaged RMSE of $\hat{\mu}$. Panel A: $N(5, 1^2)$; Panel B: $N(50, 17^2)$. Within each panel, columns are grouped by sample-size range, and each range is further split by scenario ($S_1, S_2, S_3$). Bold indicates the best-performing method within each (sample-size group, scenario) column, evaluated independently within each panel.

| | 5–10 | | | 10–30 | | | 30–50 | | | 50–100 | | | 100–200 | | |
|---|---|---|---|---|---|---|---|---|---|---|---|---|---|---|---|
| Method | $S_1$ | $S_2$ | $S_3$ | $S_1$ | $S_2$ | $S_3$ | $S_1$ | $S_2$ | $S_3$ | $S_1$ | $S_2$ | $S_3$ | $S_1$ | $S_2$ | $S_3$ |
| *Panel A:* $N(5, 1^2)$ | | | | | | | | | | | | | | | |
| BOSE | **1.10** | **1.12** | **1.02** | **1.25** | **1.16** | **1.06** | **1.35** | **1.16** | **1.09** | **1.41** | **1.16** | **1.10** | **1.48** | **1.17** | **1.12** |
| LW | 1.12 | 1.17 | 1.04 | 1.27 | 1.19 | 1.08 | 1.36 | 1.18 | 1.10 | 1.42 | 1.17 | 1.11 | 1.48 | 1.17 | 1.13 |
| BLUE | 1.21 | 1.33 | 1.13 | 1.32 | 1.29 | 1.14 | 1.40 | 1.22 | 1.13 | 1.44 | 1.19 | 1.13 | 1.49 | 1.18 | 1.14 |
| BC | 1.12 | 1.26 | 1.04 | 1.28 | 1.36 | 1.10 | 1.40 | 1.49 | 1.14 | 1.49 | 1.68 | 1.19 | 1.64 | 2.03 | 1.29 |
| QE | 1.19 | 1.15 | 1.05 | 1.50 | 1.21 | 1.15 | 1.75 | 1.26 | 1.23 | 2.05 | 1.34 | 1.32 | 2.75 | 1.51 | 1.52 |
| MLN | 1.24 | 1.24 | 1.13 | 1.36 | 1.22 | 1.13 | 1.40 | 1.26 | 1.11 | 1.45 | 1.40 | 1.14 | 1.61 | 1.76 | 1.25 |
| *Panel B:* $N(50, 17^2)$ | | | | | | | | | | | | | | | |
| BOSE | **1.10** | **1.13** | **1.02** | **1.26** | **1.15** | **1.06** | 1.35 | **1.16** | **1.08** | **1.40** | **1.16** | **1.11** | **1.47** | **1.16** | **1.13** |
| LW | 1.14 | 1.18 | 1.04 | 1.28 | 1.18 | 1.08 | 1.36 | 1.17 | 1.10 | 1.41 | 1.17 | 1.12 | 1.47 | 1.16 | 1.13 |
| BLUE | 1.23 | 1.34 | 1.12 | 1.34 | 1.28 | 1.15 | 1.40 | 1.21 | 1.13 | 1.43 | 1.19 | 1.14 | 1.49 | 1.18 | 1.14 |
| BC | 1.11 | 1.55 | 1.02 | 1.27 | 1.36 | 1.07 | 1.36 | 1.46 | 1.12 | 1.43 | 1.66 | 1.18 | 1.57 | 2.00 | 1.25 |
| QE | 1.24 | 1.26 | 1.06 | 1.56 | 1.26 | 1.17 | 2.01 | 1.34 | 1.31 | 2.87 | 1.49 | 1.57 | 4.86 | 1.70 | 2.22 |
| MLN | 1.21 | 1.20 | 1.11 | 1.29 | 1.28 | 1.07 | **1.34** | 1.51 | 1.10 | 1.45 | 2.01 | 1.24 | 1.68 | 3.15 | 1.44 |

Table S2: Averaged RMSE of $\hat{\sigma}$. Panel A: $N(5, 1^2)$; Panel B: $N(50, 17^2)$. Within each panel, columns are grouped by sample-size range, and each range is further split by scenario ($S_1, S_2, S_3$). Bold indicates the best-performing method within each (sample-size group, scenario) column, evaluated independently within each panel.

| | 5–10 | | | 10–30 | | | 30–50 | | | 50–100 | | | 100–200 | | |
|---|---|---|---|---|---|---|---|---|---|---|---|---|---|---|---|
| Method | $S_1$ | $S_2$ | $S_3$ | $S_1$ | $S_2$ | $S_3$ | $S_1$ | $S_2$ | $S_3$ | $S_1$ | $S_2$ | $S_3$ | $S_1$ | $S_2$ | $S_3$ |
| *Panel A:* $N(5, 1^2)$ | | | | | | | | | | | | | | | |
| BOSE | 1.07 | **1.90** | **0.99** | **1.42** | **2.34** | **1.13** | **1.83** | **2.54** | **1.29** | **2.45** | **2.62** | **1.44** | **3.50** | **2.68** | **1.66** |
| LW | 1.16 | 3.44 | 1.64 | 1.48 | 3.13 | 1.46 | 1.89 | 2.94 | 1.43 | 2.54 | 2.84 | 1.51 | 3.66 | 2.78 | 1.76 |
| BLUE | 1.18 | 2.28 | 1.09 | 1.47 | 2.52 | 1.17 | 1.88 | 2.64 | 1.31 | 2.51 | 2.68 | 1.46 | 3.62 | 2.70 | 1.68 |
| Shi | — | — | 1.16 | — | — | 1.24 | — | — | 1.37 | — | — | 1.51 | — | — | 1.72 |
| BC | **1.04** | 2.94 | 1.57 | 1.42 | 2.62 | 1.48 | 1.86 | 2.62 | 1.54 | 2.51 | 2.68 | 1.63 | 3.64 | 2.90 | 1.87 |
| QE | 3.41 | 2.40 | 2.25 | 2.74 | 2.67 | 2.23 | 3.20 | 2.74 | 2.65 | 4.21 | 2.75 | 3.41 | 6.19 | 2.79 | 5.07 |
| MLN | 1.40 | 2.13 | 1.27 | 2.28 | 2.63 | 1.72 | 2.77 | 2.91 | 1.97 | 3.23 | 3.08 | 2.01 | 4.25 | 3.29 | 2.04 |
| *Panel B:* $N(50, 17^2)$ | | | | | | | | | | | | | | | |
| BOSE | **1.07** | **1.96** | **0.99** | **1.43** | **2.36** | **1.12** | 1.85 | **2.60** | **1.29** | 2.41 | **2.63** | **1.44** | **3.47** | **2.65** | **1.66** |
| LW | 1.16 | 3.69 | 1.67 | 1.49 | 3.16 | 1.42 | 1.91 | 2.98 | 1.43 | 2.50 | 2.83 | 1.52 | 3.63 | 2.75 | 1.76 |
| BLUE | 1.17 | 2.41 | 1.10 | 1.49 | 2.53 | 1.15 | 1.89 | 2.69 | 1.31 | 2.48 | 2.68 | 1.47 | 3.59 | 2.67 | 1.69 |
| Shi | — | — | 1.17 | — | — | 1.21 | — | — | 1.36 | — | — | 1.52 | — | — | 1.73 |
| BC | 1.20 | 12.90 | 2.35 | 1.47 | 4.44 | 1.93 | **1.82** | 4.87 | 1.99 | **2.38** | 5.50 | 1.79 | 3.50 | 6.25 | 1.85 |
| QE | 4.55 | 20.73 | 3.28 | 2.88 | 3.56 | 2.30 | 3.19 | 3.58 | 2.63 | 3.93 | 3.55 | 3.37 | 5.49 | 3.64 | 4.73 |
| MLN | 8.42 | 3.34 | 8.58 | 8.04 | 3.79 | 6.52 | 6.67 | 4.48 | 6.04 | 4.59 | 5.08 | 4.59 | 4.12 | 6.48 | 2.82 |

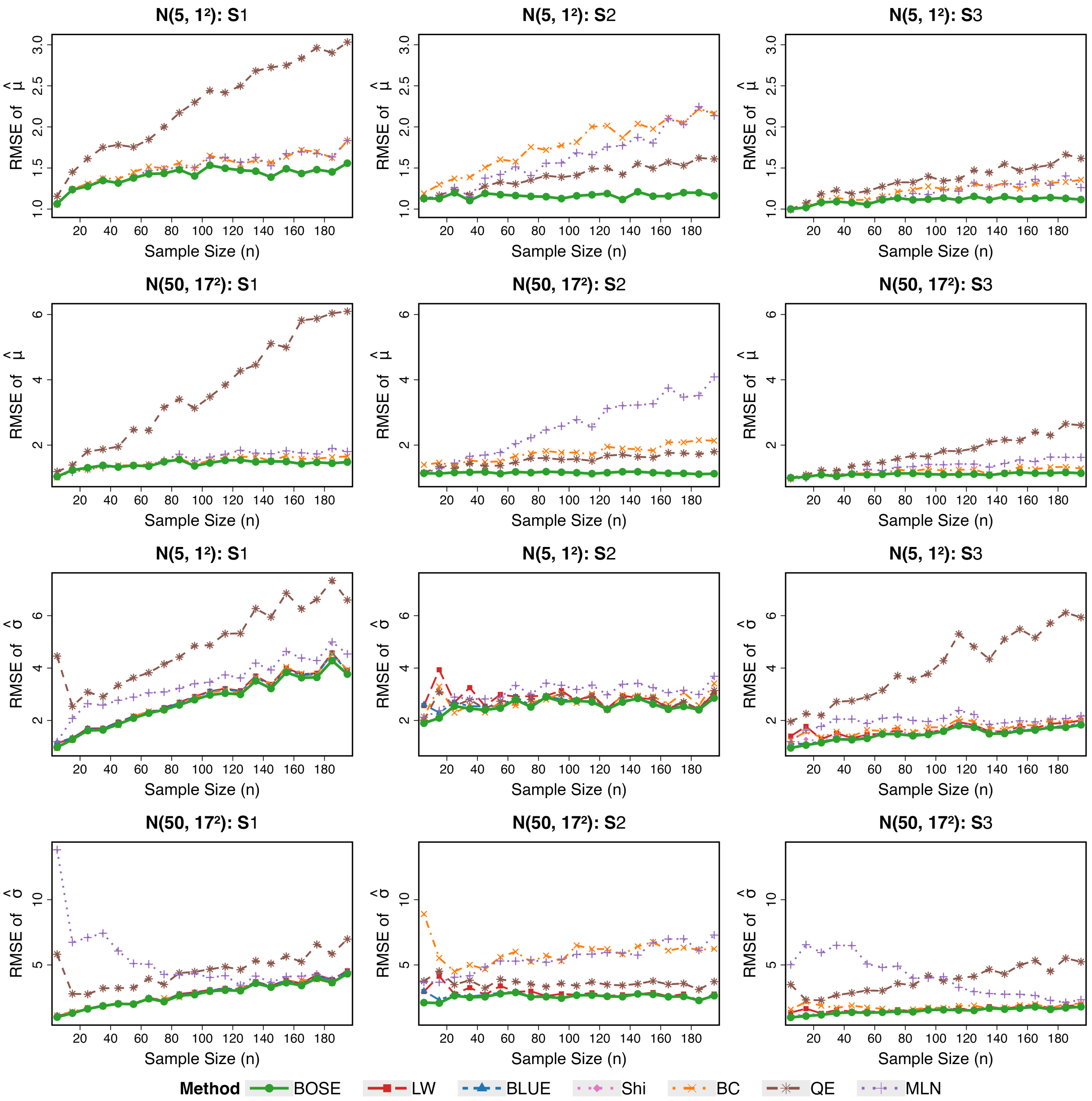


Figure S3: RMSE for mean estimation (top two rows) and SD estimation (bottom two rows) reported at individual sample sizes, using data generated from normal distributions under three common data reporting scenarios $S_1$–$S_3$. Simulations were run for every integer sample size from n = 5 to 200; results are displayed at every 10th value of n for readability.

### 4 Empirical Coverage of Credible Intervals under Normality

We evaluated the empirical coverage of BOSE's credible intervals under the normality assumption from three perspectives. First, Figure S4, presents the empirical coverage rates for the 95%, 90%, and 80% central posterior intervals given by BOSE; coverage probabilities consistently align with their nominal levels across all three data reporting scenarios ($S_1, S_2$ and $S_3$). Second, Figure S5 compares the empirical 95% coverage rates of BOSE and BLUE across sample sizes ($n \in [5, 200]$); while both methods converge asymptotically, BLUE suffers from under-coverage in small samples ($n < 50$), often dropping below 0.90 (especially under scenario $S_2$), whereas BOSE's coverage remains closer to the 0.95 nominal level throughout. Third, Figure S6 examines whether the choice of sample quantile type affects coverage: results show that coverage remains close to the nominal level across all nine quantile types (Types 1–9), with median coverage consistently within 0.94–0.95 regardless of the type used. Together, these results confirm that BOSE's credible intervals are well calibrated under normality and robust to the choice of sample quantile definition.

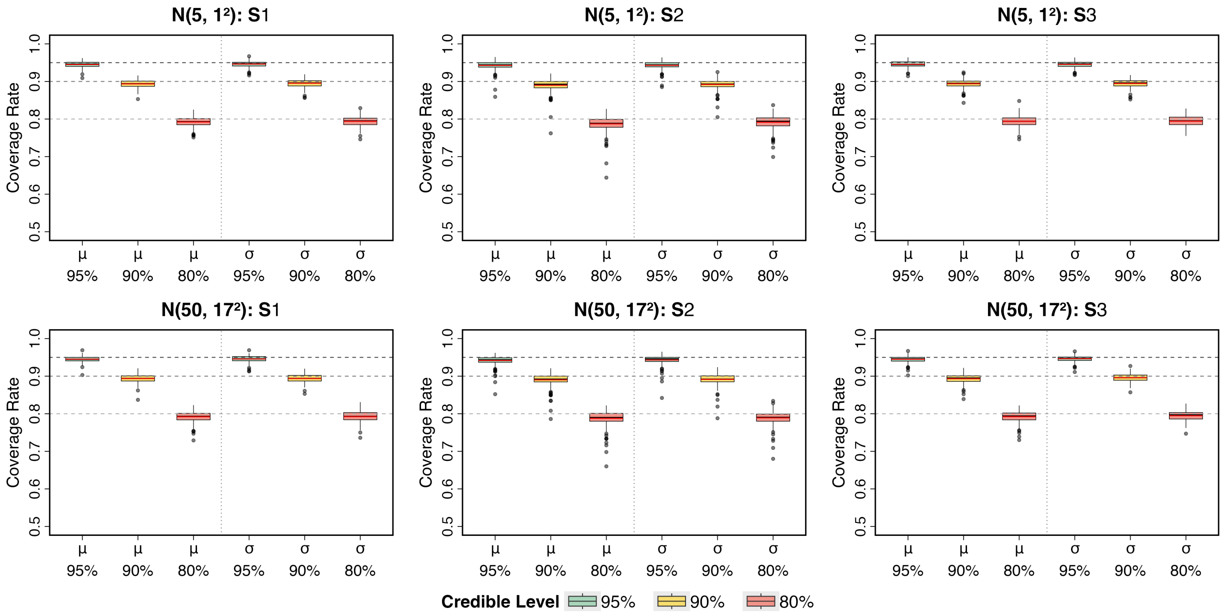


Figure S4: Boxplots of empirical coverage rates across sample sizes 5-200 for interval estimation of the mean $\mu$ and the SD $\sigma$ using 95%, 90% and 80% central posterior intervals from BOSE. Data were generated from normal distributions $N(5, 1^2)$ (top row) and $N(50, 17^2)$ (bottom row) under three common data reporting scenarios $S_1 - S_3$. The horizontal dashed lines represent the respective nominal coverage levels (0.95, 0.90, and 0.80), and the red lines within the boxplots indicate the mean coverage rates.

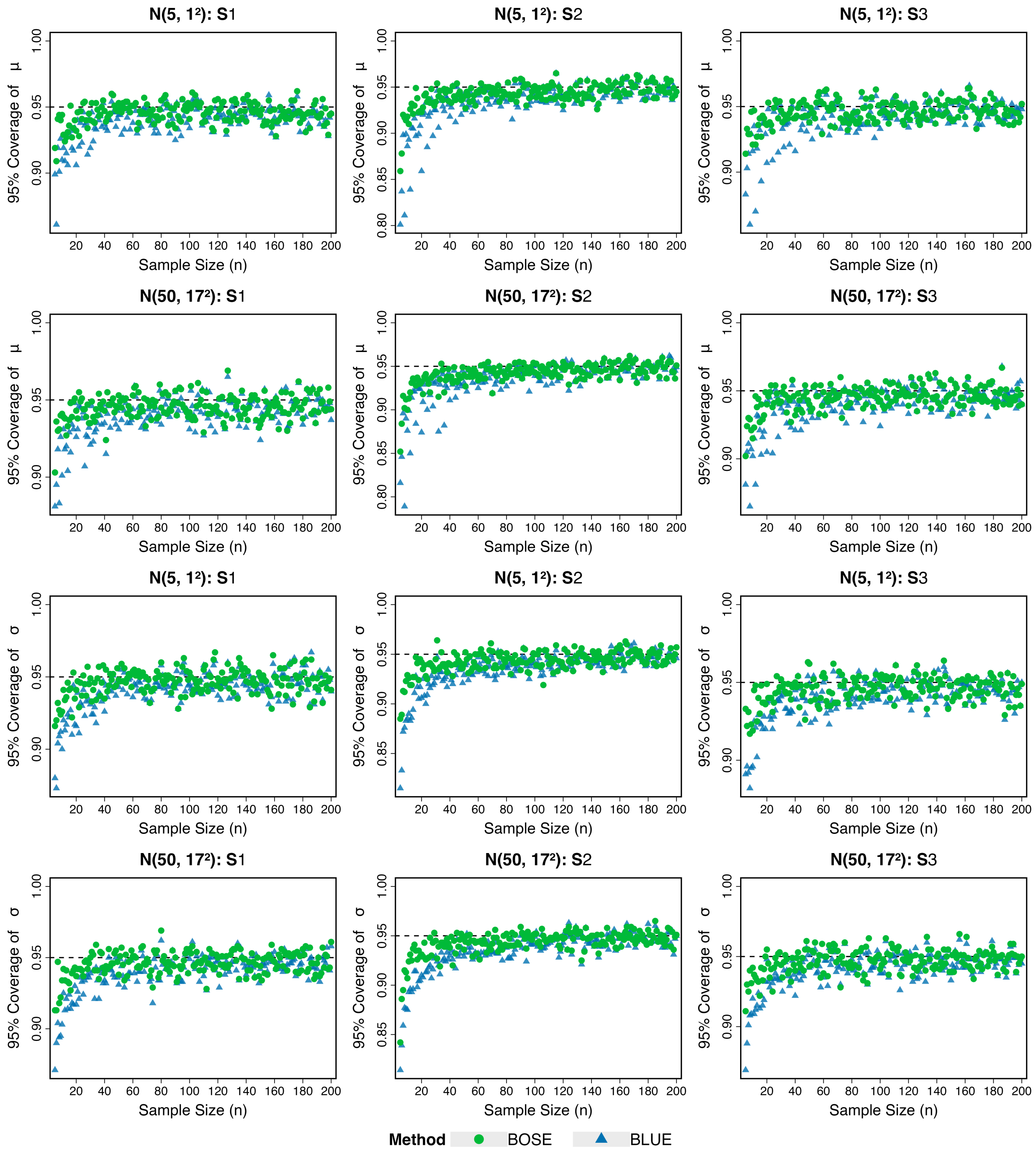


Figure S5: Performance comparison of BOSE (green dot) vs. BLUE (blue triangle) based on scatter plots of empirical coverage rates vs. sample size for interval estimation of the mean $\mu$ (top two rows) and the SD $\sigma$ (bottom two rows), using data generated from normal distributions under three common data reporting scenarios $S_1 - S_3$.

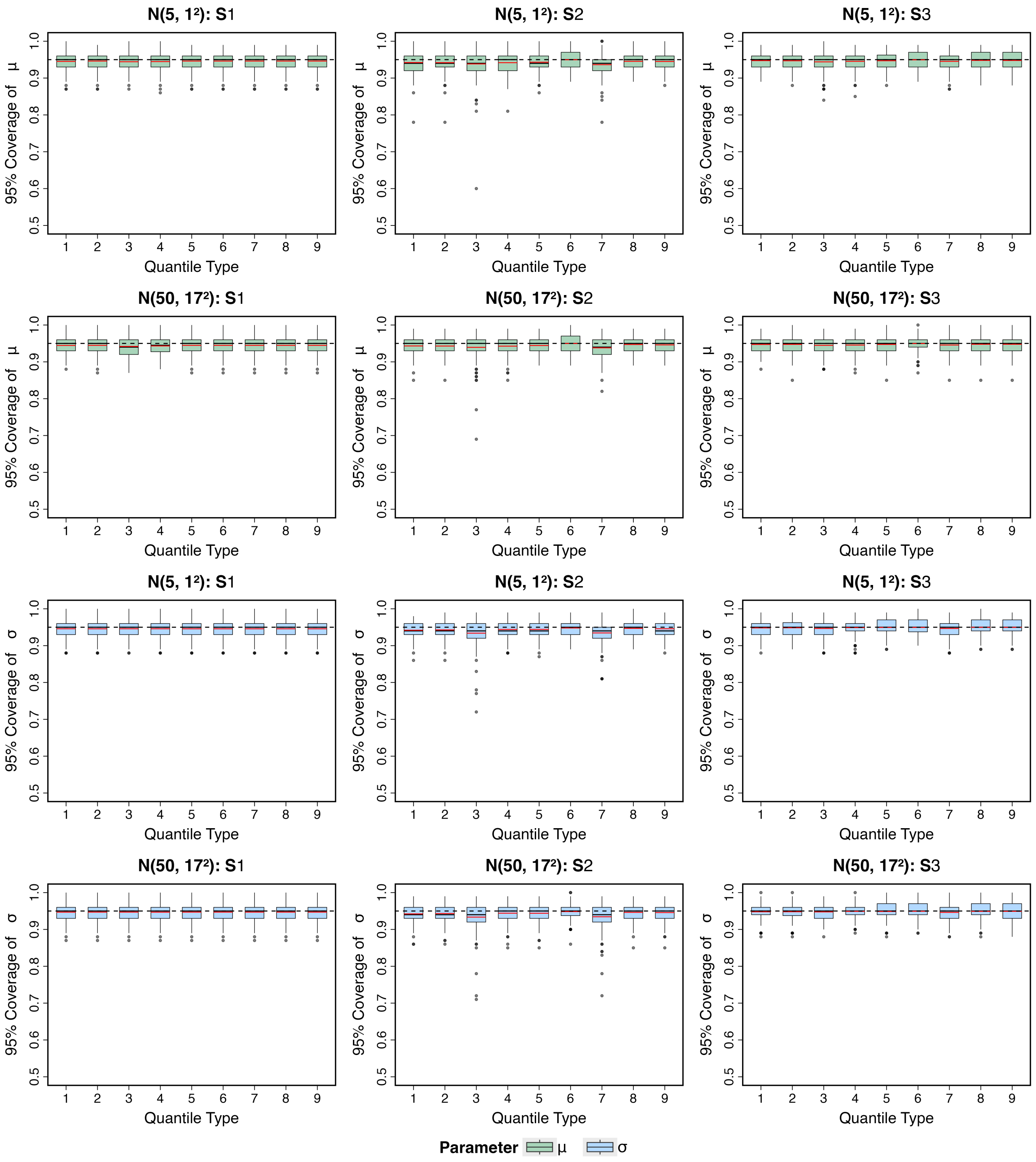


Figure S6: Boxplots of empirical 95% coverage rates for $\mu$ (top two rows) and $\sigma$ (bottom two rows) across nine sample quantile types (Types 1–9), using data generated from normal distributions $N(5, 1^2)$ and $N(50, 17^2)$ under three common data reporting scenarios $S_1$–$S_3$. The horizontal dashed line represents the nominal 95% coverage level.

### 5 Sensitivity Analysis under Extreme Non-Normality

To characterize the boundary of BOSE's applicability beyond the primary non-normal settings considered in the main text, we additionally examined performance under more extreme skewness (target skewness $\kappa \in \{1, 2, 3\}$ for the Log-Normal and Gamma families) and more extreme heavy tails (Student's t distributions with $df \in \{3, 4, 5\}$), using the same small-sample range (n = 5–50). Figures S7 and S8 report the resulting averaged RMSE for mean and SD estimation, respectively. Annotations in each panel indicate the percentage of replicates excluded due to computational failure or RMSE exceeding the threshold of 50.

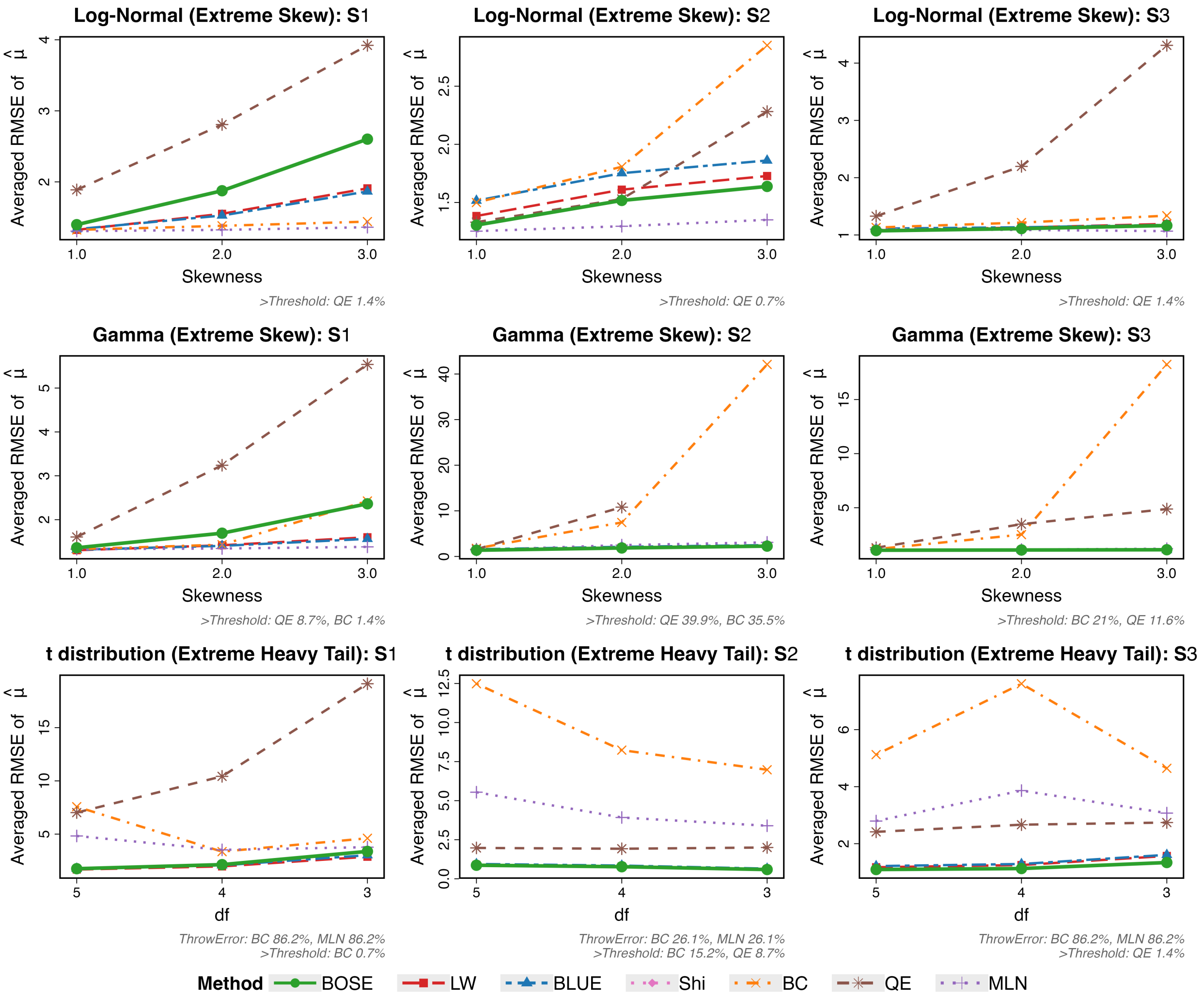


Figure S7: Sensitivity analysis under extreme skewness (Log-Normal and Gamma) and extreme heavy tails (Student's t) based on averaged RMSE for mean estimation, using data generated across three common data reporting scenarios $S_1$–$S_3$ and sample sizes 5–50. Annotations indicate the percentage of replicates excluded due to computational failure (ThrowError) or RMSE exceeding the threshold of 50 (>Threshold) for BC, QE, and MLN.

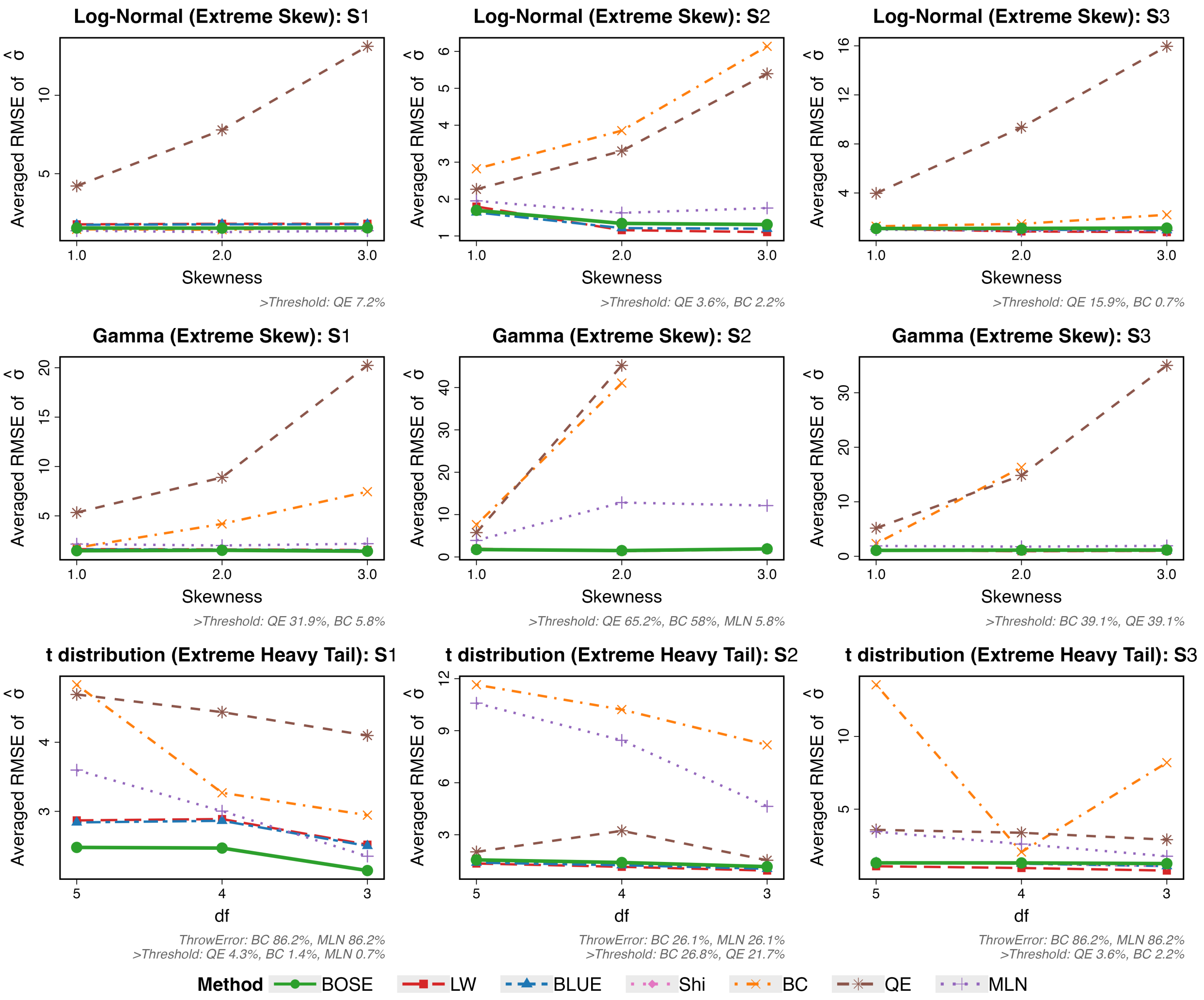


Figure S8: Sensitivity analysis under extreme skewness (Log-Normal and Gamma) and extreme heavy tails (Student's t) based on averaged RMSE for SD estimation, using data generated across three common data reporting scenarios $S_1$–$S_3$ and sample sizes 5–50. Annotations indicate the percentage of replicates excluded due to computational failure (ThrowError) or RMSE exceeding the threshold of 50 (>Threshold) for BC, QE, and MLN.

### 6 Empirical Coverage of Credible Intervals under Non-Normality

We further evaluated the empirical coverage of BOSE's credible intervals under the four non-normal distribution families considered in Section 3.3 of the main text (Log-Normal, Gamma, Student's t, and Mixture Normal). Figure S9 and Figure S10 present the resulting 95% coverage rates for $\mu$ and $\sigma$, respectively, alongside the corresponding confidence intervals from BLUE for comparison.

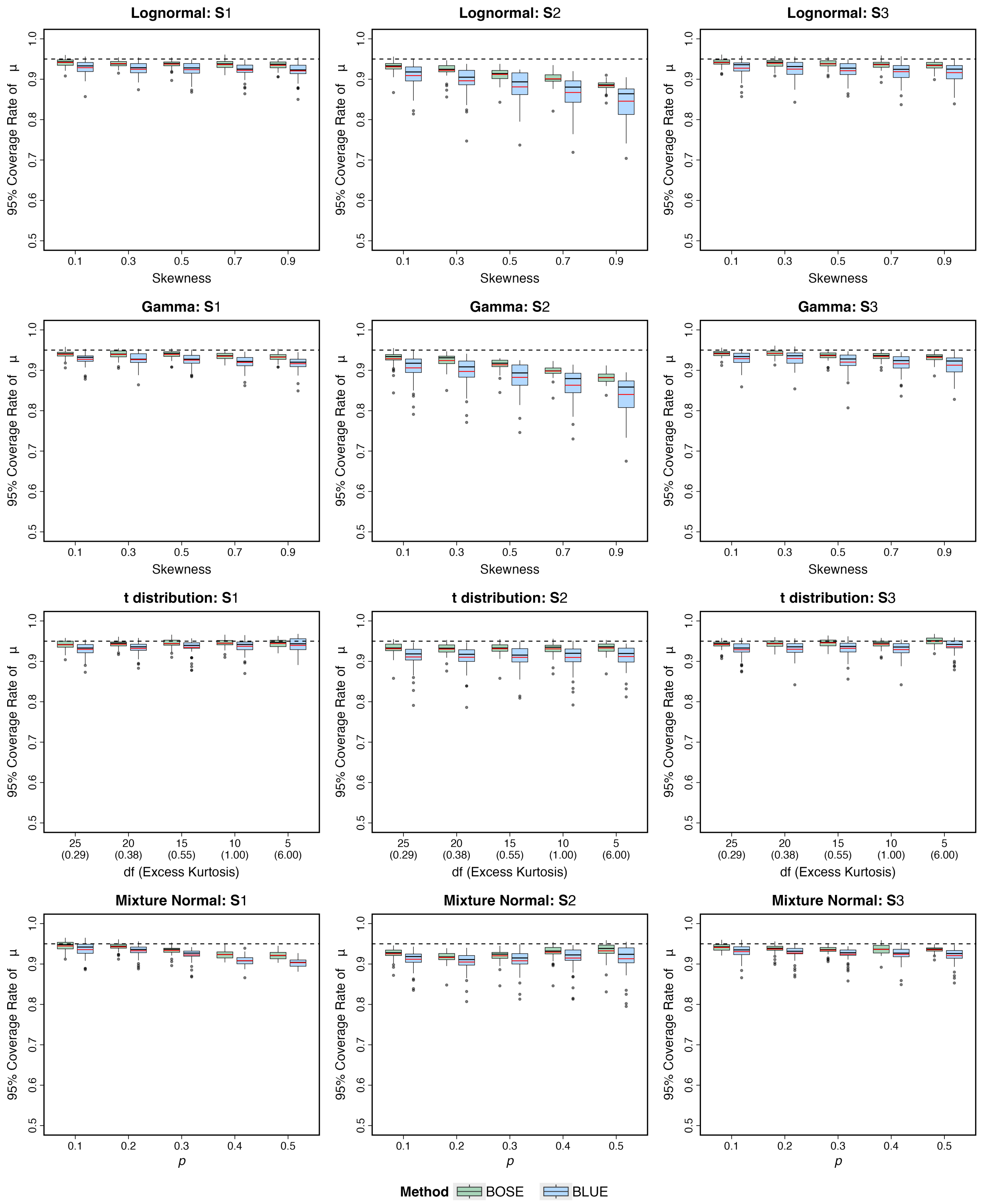


Figure S9: Empirical coverage rates of nominal 95% intervals for $\mu$ under the primary simulation settings with mild to moderate deviations from normality. BOSE credible intervals are compared with BLUE confidence intervals across four distribution families (log-normal, gamma, Student's t, and normal mixture) and reporting scenarios $S_1$–$S_3$. Each boxplot summarizes coverage rates across the evaluated sample sizes from 5 to 50, with coverage calculated over simulation replicates at each sample size. The horizontal dashed line marks the nominal 95% level; coverage closer to this line indicates better calibration.

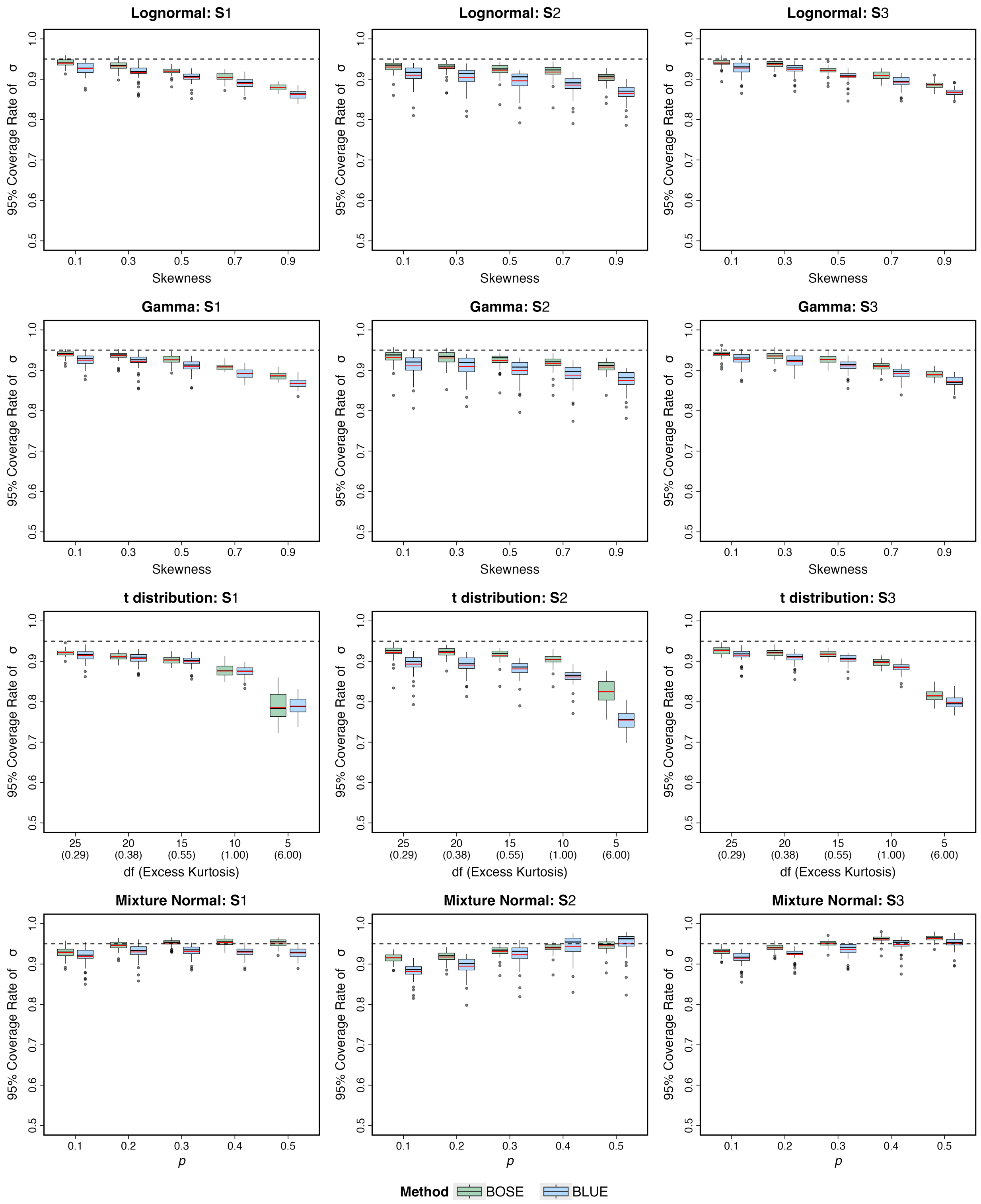


Figure S10: Empirical coverage rates of nominal 95% intervals for $\sigma$ under the primary simulation settings with mild to moderate deviations from normality. BOSE credible intervals are compared with BLUE confidence intervals across four distribution families (log-normal, gamma, Student's t, and normal mixture) and reporting scenarios $S_1$–$S_3$. Each boxplot summarizes coverage rates across the evaluated sample sizes from 5 to 50, with coverage calculated over simulation replicates at each sample size. The horizontal dashed line marks the nominal 95% level; coverage closer to this line indicates better calibration.

### 7 Data Description

Table S3: Data Example I: summary of included studies in a systematic review that examines the association between serum vitamin D levels and tuberculosis risk.

| Index | Study | Cases Size | Controls Size | Results (Serum Vitamin D Levels, nmol/L) |
|---|---|---|---|---|
| 1 | Davies, Brown, and Woodhead[1] | 40 | 40 | **Median (range):**<br>Cases: 16.0 (2.25–74.25)<br>Controls: 27.25 (9.0–132.5) |
| 2 | Grange et al.[2] | 40 | 38 | **Median (range):**<br>Cases: 65.75 (43.75–130.5)<br>Controls: 69.5 (48.5–125) |
| 3 | Davies, Grange, and Fox[3] | 15 | 15 | **Median (range):**<br>Cases: 39.75 (16.75–89.25)<br>Controls: 65.5 (26.25–114.75) |
| 4 | Davies, Brown, and Woodhead[4] | 51 | 51 | **Mean (SD):**<br>Cases: 69.5 (24.5)<br>Controls: 95.5 (29.25) |
| 5 | Chan et al.[5] | 22 | 23 | **Mean (SD):**<br>Cases: 46.5 (18.5)<br>Controls: 52.25 (15.75) |

Table S4: Data Example II: summary of included studies in a meta-analysis that examines the impact of COVID-19 on liver dysfunction.

| Index | Study | NonSurvivors Size | Survivors Size | Results |
|---|---|---|---|---|
| 1 | Chen et al.[6] | 113 | 161 | **Median (IQR):**<br>Non-survivors: 28 (18-47)<br>Survivors: 20 (14.8-32) |
| 2 | Du et al.[7] | 21 | 158 | **Median (IQR):**<br>Non-survivors: 27 (20-37)<br>Survivors: 22 (14-40.5) |
| 3 | Wang et al.[8] | 65 | 274 | **Median (IQR):**<br>Non-survivors: 24 (19-49)<br>Survivors: 28 (17-43) |
| 4 | Zhou et al.[9] | 54 | 135 | **Median (IQR):**<br>Non-survivors: 40 (24-51)<br>Survivors: 27 (15-40) |
